# The adaptive state determines the impact of mutations on evolving populations

**DOI:** 10.1101/2024.12.11.627972

**Authors:** Malgorzata Tyczynska Weh, Pragya Kumar, Viktoriya Marusyk, Andriy Marusyk, David Basanta

**Author notes:** Corresponding Author: David Basanta, Andriy Marusyk. Equal Contribution (Andriy Marusyk, David Basanta). Author contributions: Conceptualization: MTW, AM, DB, Methodology: MTW, PK, VM. Investigation and formal analysis: MTW, PK, AM, DB. Writing: MTW, PK, AM, DB. Supervision: AM, DB. Funding: AM, DB.

## Abstract

Darwinian evolution results from an interplay between stochastic diversification of heritable phenotypes, impacting the chance of survival and reproduction, and fitness-based selection. The ability of populations to evolve and adapt to environmental changes depends on rates of mutational diversification and the distribution of fitness effects of random mutations. In turn, the distribution of fitness effects of stochastic mutations can be expected to depend on the adaptive state of a population. To systematically study the impact of the interplay between the adaptive state of a population on the ability of asexual populations to adapt, we used a spatial agent-based model of a neoplastic population adapting to a selection pressure of continuous exposure to targeted therapy. We found favorable mutations were overrepresented at the extinction bottleneck but depleted at the adaptive peak. The model-based predictions were tested using an experimental cancer model of an evolution of resistance to a targeted therapy. Consistent with the model’s prediction, we found that enhancement of the mutation rate was highly beneficial under therapy but moderately detrimental under the baseline conditions. Our results highlight the importance of considering population fitness in evaluating the fitness distribution of random mutations and support the potential therapeutic utility of restricting mutational variability.

**SIGNIFICANCE STATEMENT:** The ability of a population to adapt and evolve is heavily influenced by the effects of random mutations on individuals. However, these effects can vary depending on the existing fitness level of the population. Using the development of cancer treatment resistance as an example, our research shows that populations nearing extinction can benefit from an increased rate of mutation. In contrast, mutations have a neutral or harmful effect on well-adapted populations. These findings suggest that new therapeutic strategies that manipulate mutation rates based on a population’s current state of adaptation could be effective in preventing cancer and antimicrobial resistance.

## INTRODUCTION

Populations adapt to environmental selection pressures through natural selection for phenotypic variants with higher heritable fitness. Therefore, mutational diversification, the main process responsible for the generation of heritable phenotypic diversity, is essential for creating the substrate on which selection can operate. For asexual unicellular organisms or neoplastic populations of cancer cells, a random mutation can lead to three potential outcomes at the individual cell level: fitness increase, reduction, or no change (1–3). The distribution of these fitness effects (DFE), i.e., the pool of mutations in a population before selection, can thus impact populations’ adaptation potential. This impact can be further influenced by the adaptive state of the population, which can be conceptualized using the adaptive landscape metaphor proposed by Sewall Wright (4). When a population is well-adapted, i.e., positioned at the peak of the adaptive landscape, the selective advantage of adaptive mutations is negligible, enhancing the contribution of negative mutations (Fig. 1). Thus, higher mutation rates might reduce population fitness (5). Conversely, when a population is near extinction owing to a rapid environmental change or an introduction of newly imposed selection pressure, increasing the mutation rate should increase the supply of adaptive mutations and thus be evolutionarily favorable. In the context of low fitness, deleterious mutations are still important. Indeed, critically high mutational frequencies can lead to population extinction (mutational meltdown) (6–8) for poorly adapted and well-adapted populations. Still, the supply of adaptive mutations might be more important than the occurrence of negative ones (Fig. 1). Empirical inferences of the DFEs suggest mostly negative (9–14) or neutral (15–18) fitness effects of new mutations. However, these inferences have been largely limited to well-adapted populations, thus leading to a potential bias. The dependencies of the mutational impact on the adaptive state of the population have been previously hypothesized (19, 20) but, to the best of our knowledge, not systematically interrogated.

**Figure 1.**
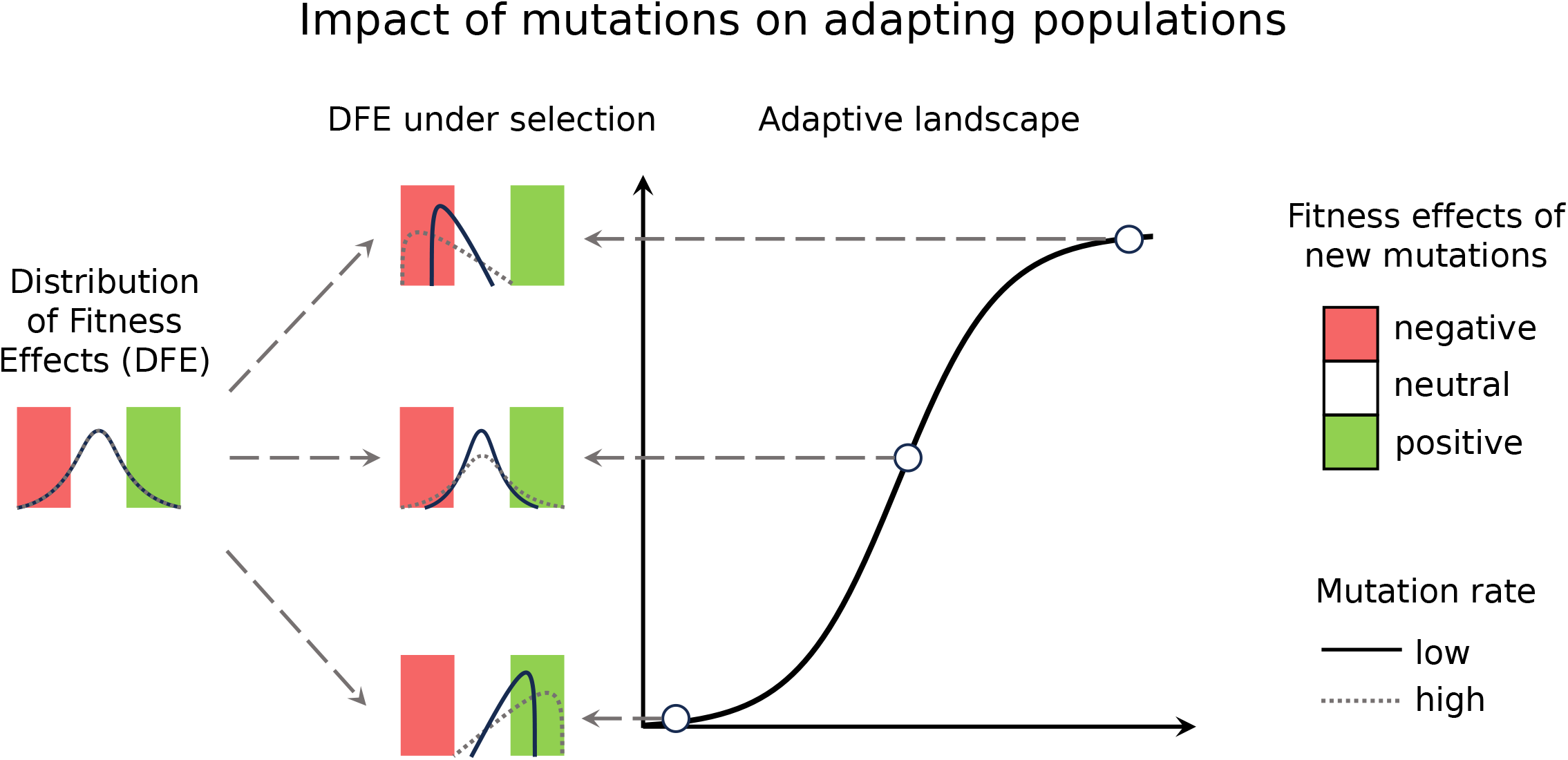
the hypothetical effect of mutations on adapting populations. The pool of mutations generated by the DFE is subject to selection, and the population’s adaptive state can determine whether this selection results in positive, neutral, or negative changes. At the top of the adaptive landscape, mutations can stabilize or decrease population fitness. At the bottom, mutations can only benefit the populations. Mutation rates emphasize these effects. Consequently, a low mutation rate can be more beneficial for the population at the top of the adaptive landscape, whereas a high mutation rate –– at the bottom.

To assess the interplay between mutation rates, DFE, and population adaptability, we used Agent-Based Modeling (ABM), a stochastic model of individual cells obeying specific actions and resulting in emergent collective behavior. In our simulations, we define a mutation as any random heritable change that can impact an individual’s fitness. This definition covers genetic alterations as well as a subset of epigenetic changes. Since space availability represents one of the key ecological resources that can substantially impact evolution, we chose to study the interplay with spatial ABM, where agents are restricted to a 2D grid domain.

Rather than studying a highly abstract population, we decided to focus on a specific scenario of a neoplastic population that faces a new therapeutic pressure that puts the population near an extinction threshold, such as an initiation of an initially effective targeted therapy. Cancer’s adaptation to therapeutic pressures might reflect a competitive release of rare subpopulations fully resistant to therapies (21). However, a growing body of evidence, including our work, indicates that, in clinical contexts with strong and durable initial remissions, therapy resistance evolves within populations of therapy persisters (22–28), whose fitness, initially very close to the extinction threshold, can be improved with mutational changes that reduce therapeutic sensitivity.

For the mathematical analysis, we first constructed the ABM capable of simulating population adaptations due to individuals acquiring mutations with various fitness effects. Next, we applied the model to study the effects of varied mutation rates on the scenario of persister cells adapting to treatment. This showed that high mutation rates can accelerate adaptations but result in substantial fitness costs long-term. Then, we extended our analytical efforts to systematically investigate the effects of mutations on adapting populations under various mutation rates, adaptive states, and DFEs. These efforts demonstrated that the adaptive state of the population determines whether the mutations will result in adaptive evolution. The mutation rates and DFEs modulate these adaptive effects.

For experimental validation of the key inferences, we assessed the dependence of the enhancement of mutation rates on the population fitness using an experimental model of ALK+ non-small cell lung cancers, the H3122 cell line. Specifically, we contrasted the impact of chemically enhanced mutagenesis between cells grown under optimal cell culture conditions versus a near-extinction exposure to clinically relevant concentrations of different ALK inhibitors (ALKi). Consistent with the model’s predictions, we found that the chemical mutagen (EMS), commonly used in experimental studies, enhances population fitness under exposure to the ALK, while moderately reducing it under optimal growth conditions. Whereas our simulations were focused on a specific context of neoplastic adaptation, our inferences should apply to other contexts of adaptation of asexual unicellular organisms, including the adaptation of bacteria to antibiotics. Our results reveal a complex, context-dependent relationship between mutation rates and adaptation, highlighting the importance of considering a population’s adaptive state when studying its evolution.

## RESULTS

### The mathematical model captures the impact of mutations on adapting populations

To understand the impact of elevated mutation rates during adaptation to treatment, we developed an Agent-Based Model (ABM): a stochastic model of agents governing emergent population behavior over time. In our model, individual agents––cells––are represented as a grid point in a two-dimensional space (Fig. 2a) and follow a set of possible actions representing a cell cycle: the cell can die, divide, and mutate with specific probabilities (rates). These actions are ordered as follows (Fig. 2b): if a cell doesn’t die, it can mutate independently of cell replication and then, if there is available space, divide and acquire a replication-dependent mutation (Methods). Because it is unknown how much mutation rates vary during the cell cycle, we set one rate for both the replication-dependent and independent mutation (p_mut_). p_mut_ is equal and constant for all cells, but cell division (p_div_) and death (p_die_) rates are cell-specific and can change if the cell is affected by a mutation. Because our ABM is spatial, the availability of unoccupied space on the grid can affect the population’s growth, especially when population size reaches carrying capacity, i.e., the maximal number of individuals on the grid. Spatial effects and stochasticity are important for population dynamics in the absence of mutations; since these cannot impact individual cells’ division or death rates, mutations are the only source of diversification in the model.

**Figure 2.**
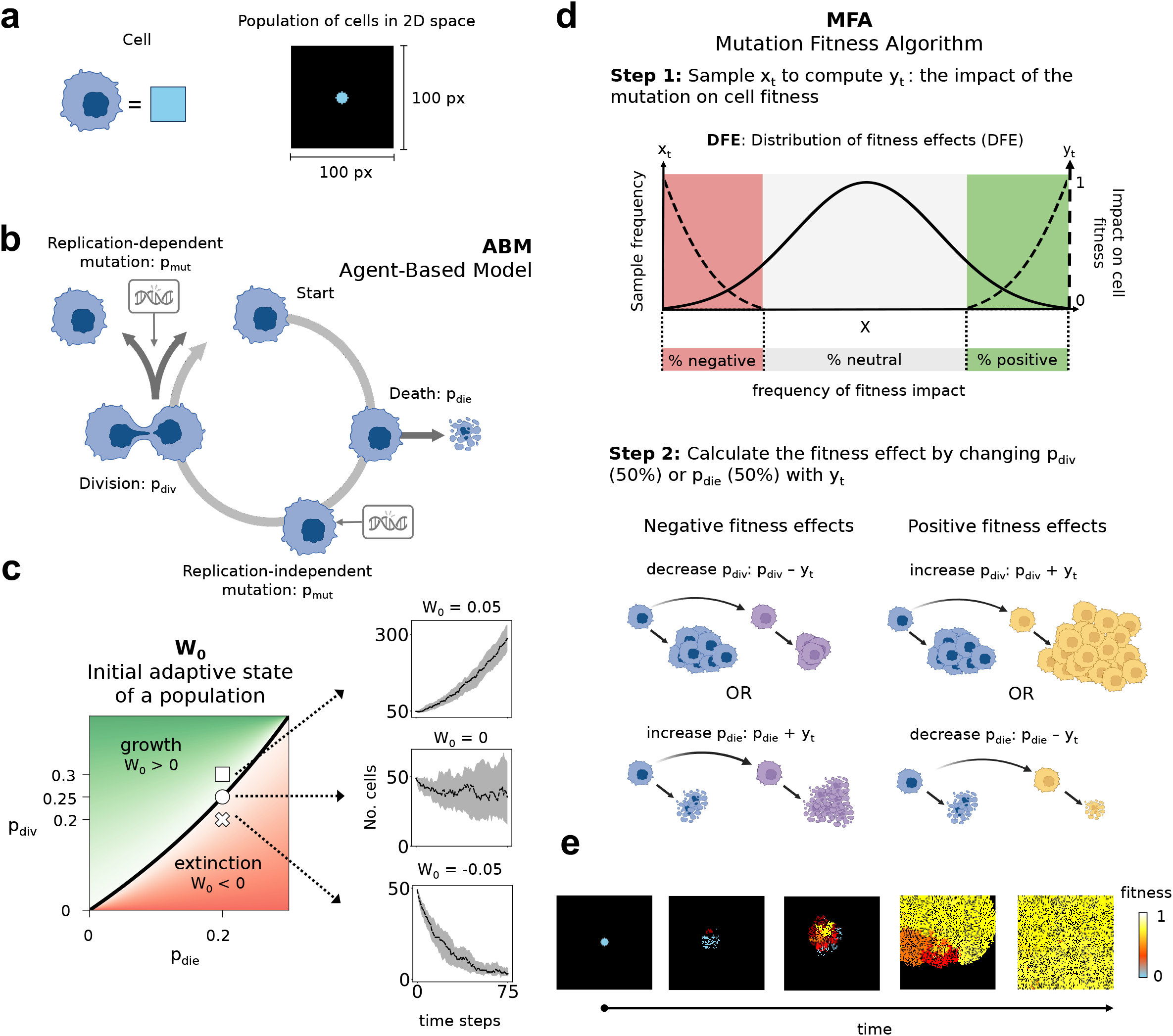
The *in silico* model to capture the impact of elevated mutation rates on adapting populations. **a**: representation of a cell as a pixel on the 100 by 100 px 2D grid. All simulations begin with cells clustered in the middle of the domain. **b**: the Agent-Based Model (ABM) flow chart. The actions are repeated for all cells. **c**: The initial adaptive state of the population (W_0_) in the p_div_, p_die_ space. The thick black line represents W_0_ = 0 and separates the (p_die_, p_div_) combinations leading to population extinction (red) or growth (green). The big, white square, circle, and cross represent W_0_ = 0.05, W_0_ = 0, and W_0_ = −0.05 values, respectively, and, when stimulated, result in growing, approximately constant, and decreasing populations (right); the gray area symbolizes 25-75% quantiles and the black line in the middle – median of 50 simulations. **d**: Visual representation of the Mutation Fitness Algorithm (MFA). Neutral mutations cause no change in fitness. **e**: snapshots of the ABM over time simulated with W_0_ = 0 and p_mut_ = 1e-1. The figure panels **a, b** and **d** are created with BioRender.com.

Next, we derived a metric to capture the initial population fitness, approximating the population’s adaptive state: W_0_ = p_div_ - p_die_/(1-p_die_). The derivation of W_0_, described in detail in Supp. Material 1, assumes that all cells are identical and well-mixed and is obtained by considering the actions of the ABM as an ordered set of conditional probabilities. W_0_ quantifies the difference between the ability to divide and the chance of death relative to survival. Since W_0_ is accurate only if all cells are identical, we initialize all cells to have equal p_div_ and p_die_ values. W_0_ = 0 results in approximately constant population over time (Fig. 2c), W_0_ < 0 leads to extinction, and W_0_ > 0 results in growth. W_0_ should not be applied to capture the adaptive state of evolved populations due to stochasticity and emergent population heterogeneity. W_0_ should also not be applied at populations near carrying capacity, as the spatial limitations can restrict population growth. Without population turnover (p_die_ = 0), W_0_ = p_div_.

In our ABM, mutations change the individual cell fitness (w ≔ p_div_ - p_die_) by affecting either division or death probability with a quantity determined by a Mutation-Fitness Algorithm (MFA). This quantity is stochastic and generated from a specific distribution of fitness effects. The MFA results in an immediate and heritable fitness change in individual cells; the algorithm is sketched in Fig. 2d and proceeds as follows (see Methods for details and Supp. Mat. 1 for mathematical analysis): First, we compute the mutation’s impact on cell fitness (Step 1 on Fig. 2d). To reflect the mutation’s stochastic nature, we sample a random number x_t_ ∼ X; depending on the value of x_t_, the impact is negative, neutral, or positive. We compute the magnitude of this impact by linearly transforming x_t_ to y_t_. Next (Fig. 2d step 2), depending on whether the mutation is positive or negative, we add or subtract the y_t_ to/from p_div_ or p_die_; the probability of mutation changing either p_div_ or p_die_ is 50%. The new values of p_div_ or p_die_ and the number of mutations are inherited during cell division. Cell fitness changes due to the calculated impact of a mutation, and cell fitness is invariant to the number of mutations a single cell has acquired.

We run a subset of initial simulations visually tracking evolution to ensure that our ABM can capture emerging adaptation. For example, the snapshots from a simulation with p_mut_ = 1e-3 (Fig. 2e) demonstrate population-level adaptations emerging from the gradual fitness increase. In addition, the set of Supplementary Movies S1-S6 shows that the ABM model can capture either population extinction or adaptation when subject to elevated mutation rates. Hence, our ABM provides a platform for studying many evolutionary trajectories.

### At the edge of extinction, higher mutation rates facilitate adaptation at the expense of decreasing maximal population fitness

We first apply our model to investigate how mutations impact adaptations of persister cells to treatment. In our simulations, we define persister cells as a sub-population of therapy-naive cells that is capable of surviving the initial therapeutic elimination but initially has a near-zero growth rate under therapy (22–28). Populations of persisters can evolve bona fide therapy resistance through a mutation-selection process. To model the adaptation of persister cells to cancer treatment, we initialize the simulations with 49 cells in a (100 x 100) px grid, thus with a carrying capacity of 10000 cells. This proportion of persisters was chosen based on the quantitative estimates in the literature (23) (Materials and Methods). We then set their initial adaptive state to W_0_ = 0 (i.e., zero net growth) with p_div_ = 0.25 and p_die_ = 0.2. We also set p_die min_ = 0.196; this ensures a high cell turnover and is inspired by experimentally derived estimates of tumor death rates (29, 30). Because W_0_ = 0 corresponds to the fitness of persister cell populations under treatment, we do not model treatment explicitly.

While the specific frequencies of the positive, negative, and neutral mutation vary widely across the literature, and many potential DFEs can govern adaptations in persister cell populations, we decided to initialize the MFA with the DFE of a simple case of a distribution having equal, small chances of negative and positive fitness effects (15). This choice results in the frequency of neutral mutations as 99% and 0.5% of negative and 0.5% of positive mutations (see Supp. Mat. 1, Materials and Methods). We initialized each ABM with mutation rates p_mut_ = 0, or in a range 1e-4–1 (see Materials and Methods). These relatively high mutation rates aim to compensate for the relatively small population sizes in our simulations compared to the residual tumors in clinics. While our time step is abstract, we consider the unit of every time step as ½ day and scale the time steps accordingly to hypothetical years.

Consistent with the expectation of mutations enhancing the ability of populations to adapt at the edge of extinction, an increase in the mutation rate prevents population extinction (Fig. 3a) and accelerates adaptation (Fib 3b). On the other hand, the steady-state values of the average population fitness 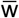 (Fig. 3c-d) and population numbers (Supp. Fig. 5a) decrease with increasing mutation rates, suggesting potential fitness costs of high mutation rates. Notably, changing the initial adaptive state of a population to a net negative growth rate (W_0_ = −0.146 with p_div_ = 0.03 and p_die_ = 0.15) or to a near-zero net growth rate but with lower division and death probabilities (W_0_ = −0.0014 with p_div_ = 0.175 and p_die_ = 0.15) does not qualitatively affect the results (Supp. Fig. 9).

**Figure 3.**
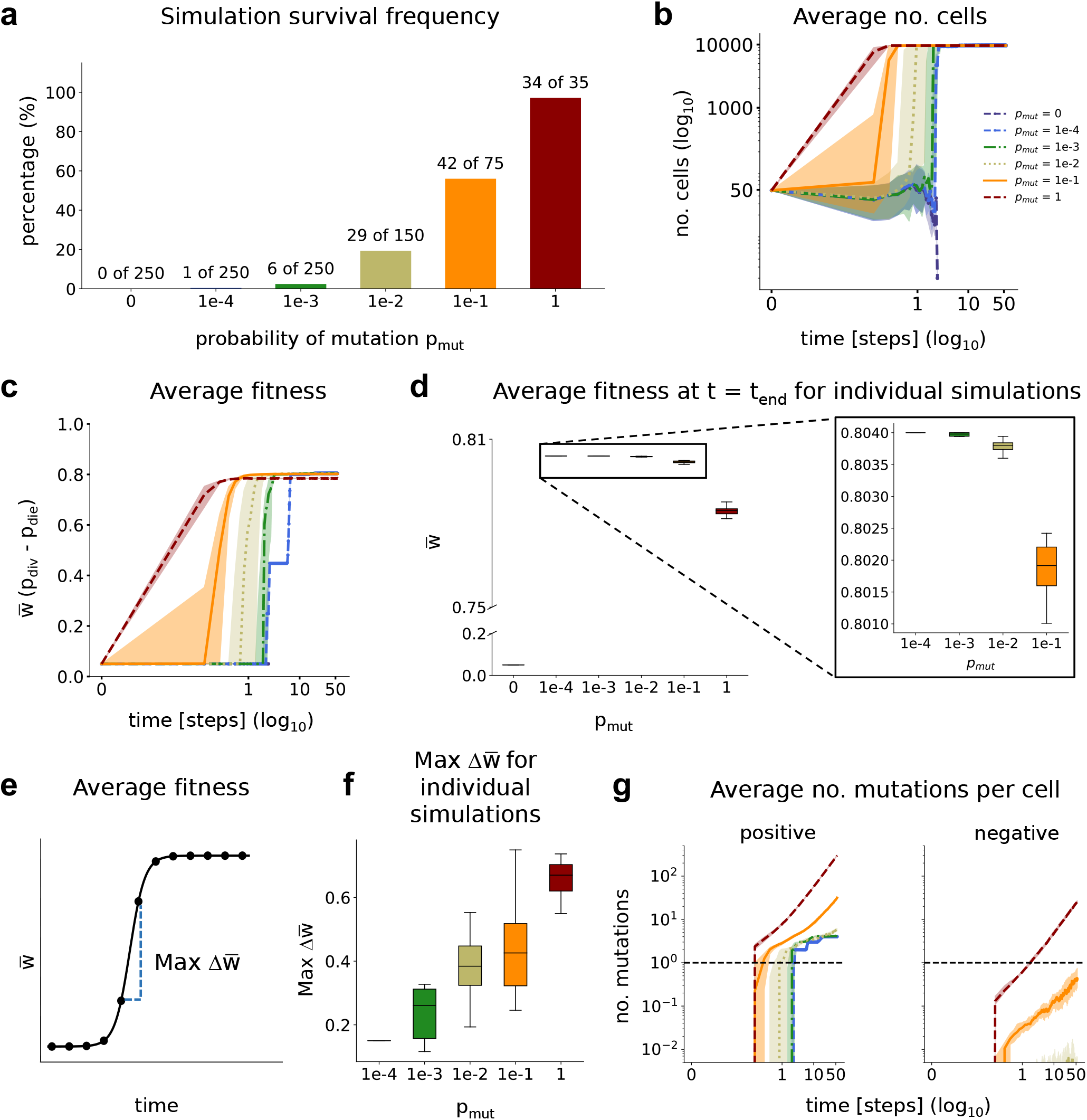
The qualitative predictions of the impact of the elevated mutation rates on the persister cell adaptations to treatment. The persister cell state under treatment is represented by a small population number (49 cells) and zero net growth rate (W_0_ = 0). **a**: the percentage of replicates surviving until the end of the simulation (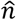; t_end_ = 40000 time steps) out of (n) simulations. **b**: the average population number over time (log_10_-log_10_). **c**: the average fitness over log10 time, and **d**: the corresponding distribution of the median average fitness at the end of the simulation (t_end_) across 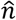 replicates (left), with a focus on the p_mut_ = 1e-4 — 1e-1 (right). The differences between distributions with 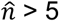 are significant on the 1% level (two-way KS statistics). **e**: the average number of positive (left) and negative (right) on the log_10_-log_10_ scale. The color scheme for all graphs follows **a. f**: Illustrative example of the Max 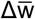. **g**: Max 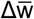 for the individual simulations over p_mut_. For **b, c**, and **e**, the thick lines represent the median average across 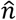 simulations, and the colored areas are the corresponding 25-75% quantiles. For all graphs, we scaled the time step unit to hypothetical years, assuming 1 time step = ½ day.

**Figure 4.**
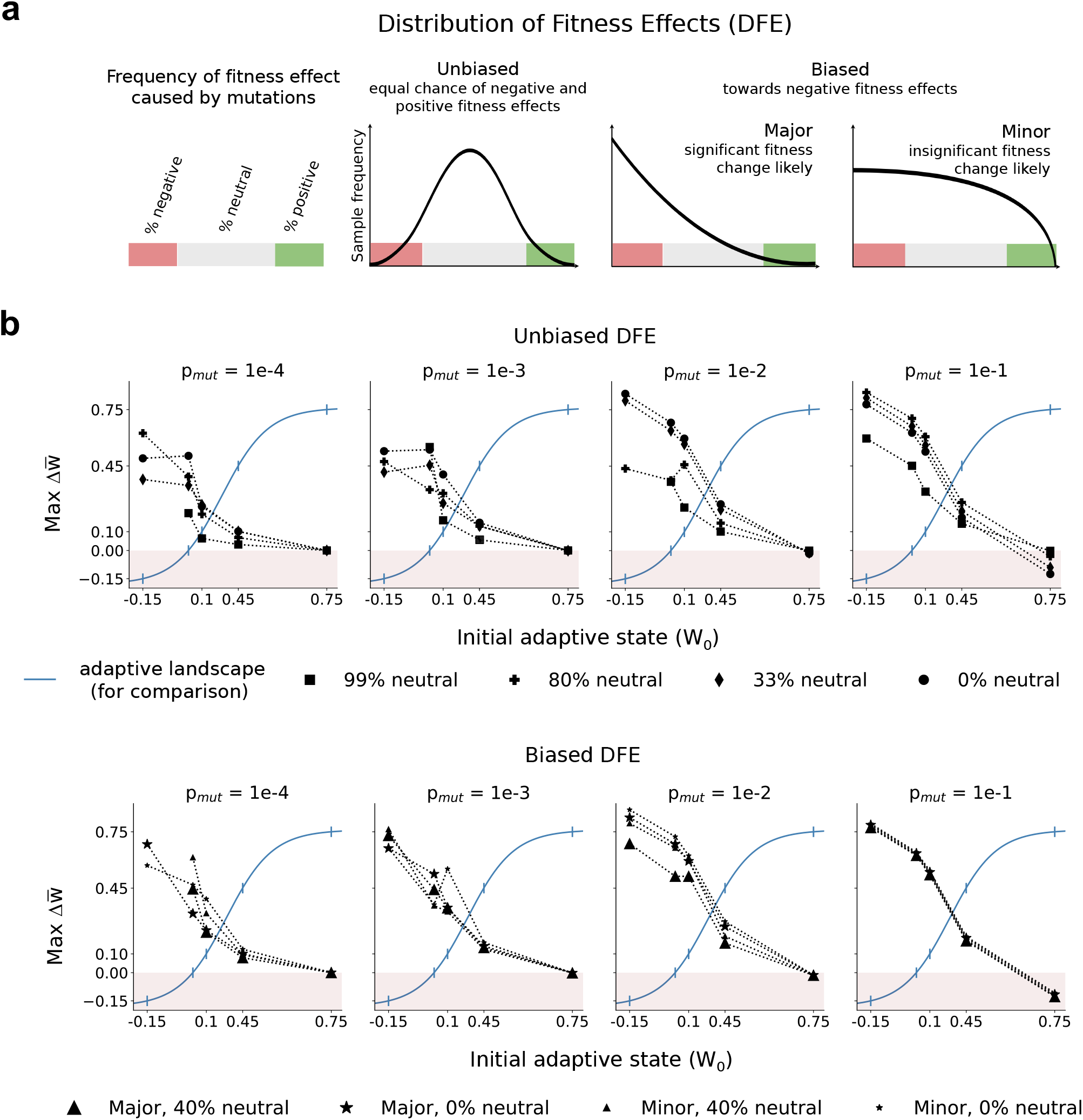
The impact of elevated mutation rates changes with the position on the adaptive landscape and the choice of the distribution of fitness effects. **a**: a visual representation of the groups of DFEs. **b**: Max 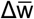 (y-axis, thick, black) of *n* simulations initialized at specific adaptive states W_0_ (x-axis). The values are displayed relative to the adaptive landscape (think, blue line) since Max 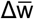 and W_0_ share the same numeric values. The black symbols represent a specific choice of the DFE and the pink region of Max 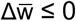 represents the initial conditions leading to extinction in the absence of adaptation. The W_0_ values for simulations are obtained with p_die_ = 0.2, and p_div_ = (0.1, 0.25, 0.35, 0.7, 1). For each combination of W_0_ and DFE, the *n* is 35 (p_mut_ = 1), 75 (p_mut_ = 1e-1), 150 (p_mut_ = 1e-2), or 250 (p_mut_ = 1e-3, 1e-4, 0). Some graphs are missing because of population extinction before the end of the simulation. The graphs for the extreme conditions (p_mut_ = 0 and 1) are displayed in Supp. Fig. 7.

**Fig. 5.**
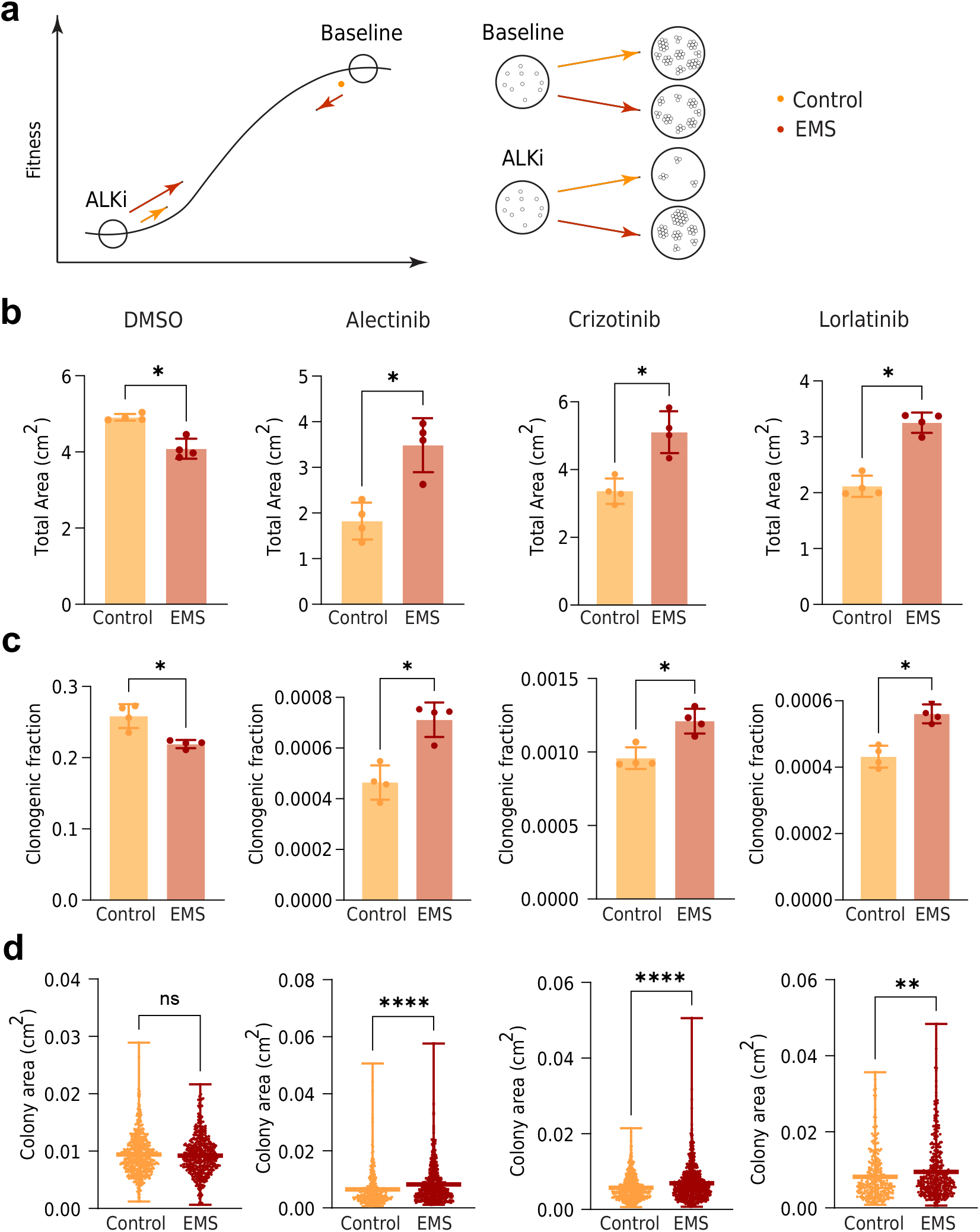
*In vitro* model of the mutagen effect on the persister cells’ adaptation to treatment. **a:** A schemata of the experimental design. H3122 cells were treated with DMSO (baseline) or ALKi, Alectinib (0.5µM), Crizotinib (0.5µM) or Lorlatinib (2µM), alone or in combination with EMS (1µM). For the EMS groups, cells were pre-treated with EMS for 48 hours. Cell colonies were analyzed. **b:** Total area covered by the cells. **c:** Clonogenic fraction (number of colonies as a fraction of cells plated). **d:** Area of each colony. For each graph, the significance testing was done with the Mann-Whitey test; p-values – <0.05 (*), <0.01 (**), <0.001 (***), <0.0001 (****).

To measure the speed and the direction of the evolution, we derived a metric Max 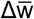: maximum average change in fitness (Fig. 3e and Methods). Max 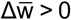 indicates fitness increase leading to adaptation, and Max 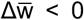––population fitness decline. As expected, Max 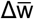 increases with the increase in mutation rate (Fig. 3f, Supp. Fig. 3b).

To investigate the causes of the fitness costs, we analyzed the number of accumulated positive and negative mutations (Fig. 3g). While we observe a substantial accumulation of positive mutations for all mutation rates, the average number of negative mutations is several orders of magnitude smaller. Despite this, populations with high mutation rates accumulate negative mutations, suggesting that the fitness costs emerge from the total load of negative mutations (Supplementary Movies S7-S18). The number of positive mutations acquired at the time of Max 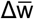 approximates how many mutations are required for adaptive evolution (Supp. Fig. 3b). Our analysis shows that regardless of the mutation rate, 1-3 mutations are necessary for adaptations, which matches the estimated number of mutational events contributing to tumor expansion (31) or the emergence of antibiotic resistance (32, 33) from the literature.

### The impact of mutations on adaptive evolution depends on the adaptive state of the population

Having validated the intuitive prediction that mutations benefit populations at the edge of extinction but reduce their fitness in the long term, we sought to investigate the impact of mutations on evolving populations systematically. Initially, we assumed a symmetric DFE distribution with rare deleterious fitness effects. This is likely wrong because prior studies suggest mostly neutral or deleterious fitness effects of new mutations (9–18). Thus, we considered a variety of distributions, which we further divided into two groups (Methods) (Fig. 4a): the first group is made of distributions with equal frequencies of positive and negative effects. Thus, it is unbiased about either negative or positive fitness effects. The second group comprises distributions biased towards deleterious fitness effects; we refer to these groups as “unbiased” and “biased.”; we have not investigated DFEs with a bias towards positive fitness effects due to the lack of such DFEs in the literature. Methods and Supp. Mat. 1 provide a mathematically precise definition of all the distributions.

In the unbiased group, lower frequencies of neutral mutations lead to an increased pool of positive and negative mutations. Because of selection, we expect an increased speed of evolution with a decreasing frequency of neutral mutations. Thus, we set the distributions from the unbiased group to have neutral fitness effects of 99%, 80%, 33%, and 0% (corresponding to 0.5%, 10%, 33%, and 50% of positive/negative effects, respectively). This wide range of frequencies reflects previously reported findings on 98-99% of neutral point mutations (15), 10% of fitness increases during adaptation (34), and neutral mutation frequencies of less than 30% (10).

The group of DFEs with a bias towards deleterious fitness effects reduces the pool of mutations leading to adaptive changes. Since the chances of successful adaptations can depend on 1) the variety of negative fitness effects and 2) the frequency of the positive and neutral mutations, we further divided the biased DFEs into two groups: one having a major bias towards very deleterious mutations (labeled as “Major”) and one less biased, allowing for a large variety of negative fitness effects (labeled as “Minor”). The distributions from the biased group always have 40% negative effects but can have either 40% neutral effects (i.e., 20% positive effects) or 0% neutral effects (i.e., 60% positive effects).

Since we speculate that the mutation’s impact on evolving populations varies with the adaptive state of the population, we also vary W_0_ (recall Fig. 2c). We keep the high population turnover by setting p_die_ = 0.2 and vary the initial p_div_ to model different degrees of adaptation (Fig. 4b): population extinction (W_0_ = −0.15), zero net growth (W_0_ = 0), intermediate population growth (W_0_ = 0.1, 0.45) and maximum possible growth, representing the top position on the adaptive landscape (W_0_ = 0.75).

We again apply the Max 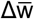 to test whether the evolution is adaptive in all 240 sets of simulations (six different p_mut_, eight different DEFs, and five W_0_ values), each initialized with 49 cells. In addition to each Max 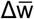 measurement, we record the trajectories of the average population sizes, fitness, number of positive/negative mutations, and more; we provide an example of illustrative cases in the Supp. Fig. 5. Finally, Methods and Supp. Fig. 4 assesses the required number of simulation replicates for statistical significance.

Fig. 4b demonstrates that the impact of mutations on adaptive evolutions varies with the adaptive state of the population. At the bottom of the adaptive landscape (W_0_ = −0.15, 0), higher mutation rates increase the odds of adaptation. For better-adapted populations (W_0_ = 0.1, 0.45), mutations still lead to adaptive evolution, but the lower magnitude of Max 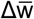 implies a less dramatic effect. At the top of the adaptive landscape (W_0_ = 0.75), frequent deleterious mutations result in maladaptive changes (Max 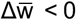); only populations with absent or infrequent mutations can maintain high initial fitness (Max 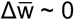), indicating the mutation-selection balance. Overall, these results indicate that mutations determine the effects of mutation based on the adaptive state of the population.

Regardless of the DFE, the graphs of Max 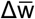 over the adaptive landscape are more aligned to the W_0_ = 0 line as the mutation rate decreases; p_mut_ = 0 all Max 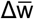 are 0 if W_0_ > 0 for all DFEs (Supp. Fig. 8). This behavior is not unexpected: a lower rate of diversification reduces the pool of mutations that the selection can act upon and, consequently –– the reduced propagation of adaptive/maladaptive effects of mutations.

Changes in the DFE can cause variation in Max 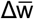 for a given mutation rate and adaptive state: for the unbiased DFE, p_mut_ = 1e-1, the Max 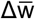 is maximized at 80% of neutral mutations for all W_0_ ≠ 0.75, and for p_mut_ = 1e-2 under the same conditions - at 0% of neutral mutations. For p_mut_ = 1e-1 and p_mut_ = 1 (Supp. Fig. 8), we observe mutational meltdowns for some, but not all forms of DFEs: for the biased DFEs at p_mut_ = 1, the meltdown happens for all DFEs except the DFE with a minor bias towards negative fitness effects and the 0% of neutral mutations. Collectively, this shows that the relationship between the DFE and the potential for adaptive evolution is nonlinear: how the specific distribution will dictate evolution is difficult to predict, even if the mutation rate and the adaptive state of the populations are known.

### Experimental studies support the predicted interplay between the adaptive state of a population and the impact of enhanced mutation rates

To empirically test the theoretical findings of the dependence of the impact of enhanced mutation rates on population fitness, we decided to contrast the effect of enhanced mutations between optimal growth conditions and therapy-induced selection bottleneck under an experimental context that closely mirrors the scenario explored in our in silico simulations. To this end, we used an experimental cancer model, the H3122 cell line, a commonly used model to study acquired therapy resistance in targetable ALK+ lung cancers. This established cell line is characterized by a stable, high growth rate under standard *in vitro* cell culture conditions. Exposure to clinically relevant ALKi concentrations eliminates the majority of therapy-naive cells; the surviving minor fraction initially displays a persistence phenotype (24). These persisters gradually adapt to the drugs by accumulating multiple resistance-promoting mutational and expression-level changes, eventually approaching the baseline growth rates of the therapy-naive H3122 cells (24). Notably, our ABM provides a reasonable approximation of this experimental system. Both in silico and in vitro, cells grow in 2D monolayer, with minimal migration; cell proliferation requires space availability. Whereas in H3122 cells, the majority of resistance-promoting changes occur via a non-mutational mechanism, the adaptation relies on some mutational changes, and expression level variability includes a substantial stochastic component (35).

Based on the in silico results, we expected that elevation of stochastic mutation rates, induced by continuous exposure to non-toxic concentrations of a chemical mutagen EMS, should enhance the speed of adaptation of therapy-exposed cells while having either no effect or cause a modest reduction of population fitness of cells under the baseline growth condition (Fig. 5a, left). To detect the impact of the enhanced mutation rates, we adopted a clonogenic assay used in our earlier study (24). Specifically, cells were seeded at clonal density in the presence of clinically relevant ALKi concentrations or DMSO (vehicle control), with and without EMS. Following 2.5 weeks (DMSO) or 3-5 weeks (ALKi) clonogenic fractions and sizes of colonies were evaluated as a fitness readout (Fig. 5a, right) (24).

Consistent with the theoretical predictions, in the absence of therapy, EMS exposure led to a slight reduction of clonogenic survival of H3122 cells and the overall population size (combined area of all colonies). While the distribution of colony sizes trended smaller under EMS treatment, the difference was not statistically significant (Fig. 5b-d, Supp. Fig. 8). As expected, treatment with three different ALKi (alectinib, lorlatinib, and crizotinib) reduced the clonogenic fraction and inhibited the growth of clonogenic cells (reflected in colony sizes), compared to the DMSO control (Figure 5b-d, Supp. Fig. 8a). At the same time, clonogenic fraction, total population size, and size of individual colonies were significantly larger under the EMS exposure, indicating enhanced adaptability of the populations under therapeutic bottlenecks (Fig. 5b-d, Supp. Fig. 8a). Importantly, EMS did not impact ALKi sensitivity in short-term viability assays (Supp. Fig. 8b), indicating that the enhanced clonogenic proportion and colony sizes reflected the EMS-enhanced evolvability rather than a direct decrease in ALKi sensitivity. In summary, experimental studies supported the in silico inferences, indicating that the impact of enhanced mutation rates depends on the state of the population fitness.

## DISCUSSION

Our computational and experimental studies demonstrate that the fitness effects of mutations are not fixed but rather shaped by a population’s adaptive state. This finding validates and extends beyond the intuitive expectation that mutation outcomes should vary based on a population’s position in the adaptive landscape. At the extinction threshold, enhanced mutation rates increase the supply of adaptive mutations enabling the evolutionary rescue (36). The importance of mutational diversification for evolutionary rescue under ecological constraints is well appreciated and has been studied in multiple theoretical models (37–40). However, once populations adapt, the positive mutations cease to provide a selective advantage, thus tipping the balance towards the effects of negative mutations and higher mutation limits the maximal fitness of a population. At the peak of the adaptive landscape, high mutation rates disrupt the mutation-selection balance, reducing population fitness, while low rates preserve balance and maintain fitness. Our findings highlight the importance of considering the adaptive state of populations when interpreting experimental DEF inferences. This dependency on the adaptive state can profoundly impact the consequences of random mutations (19–20, 41), yet it’s often overlooked. Finally, our study focused on neoplastic populations exposed to therapy, our conclusions should be generalizable to other contexts involving asexual populations, such as yeasts and microbes.

An accurate understanding of the effects of mutation rates on evolutionary dynamics requires adequate quantitative knowledge of the DFE. However, empirical inferences of DFEs are complicated by the strong dependence of observed DFE on the adaptive state of populations. Most random mutations can be expected to reduce fitness or be fitness-neutral, as disrupting complex molecular processes is easier than improving them (42). This intuition is supported by multiple empirical studies, revealing a strong bias towards neutral or mildly deleterious effects of random mutagenesis (9–18). However, intuition is unsuitable for quantitative inferences, and most experimental studies have operated with well-adapted populations near or at the fitness peak (11, 13–14, 17). This scenario facilitates well-controlled inferences but limits the ability to detect the effect of positive mutations. Similarly, computational statistical inferences such as mutational analyses of cancer genomes (9, 15, 18) are likely confounded by a reflection of relatively long periods of relative stasis with tumor expansion occurring at a local fitness plateau.

The use of spatial ABM enabled us to sidestep some of the significant limitations of both experimental and statistical inferences of DFE. ABM-based studies enable simultaneous tracking of the fitness of individuals, mutational occurrences, and their effects. This approach allows for resolution unachievable by the population-genetics-based mathematical models of fitness effects of new mutations, such as the Gillespie mutational landscape model (43) and Fisher’s geometric model (44). Importantly, our studies highlight the importance of considering space explicitly: despite having a division probability that significantly exceeds the probability of death (Fig. 2c), populations exhibited near-zero growth due to the lack of free adjacent space available for cell division. The corresponding rate of change in average population fitness is thus dependent on the carrying capacity and the population turnover rate. While the importance of space is often overlooked, space represents a critical ecological resource(8) that profoundly affects eco-evolutionary dynamics in contexts of cellular populations with relatively low motility (45, 46). Consequently, a lack of spatial considerations can lead to biased inferences and misunderstanding of the drivers of the evolutionary dynamics.

Our findings have important therapeutic implications. Our results are consistent with the phenomenon of stress-induced mutagenesis, i.e., the elevation of mutation rate under therapeutic and environmental stress with a return to baseline rate once populations have adapted to the stressor (28, 47–49) activation of APOBEC3A in lung cancers can drive the evolution of persister cells under EGFR-targeted therapy (50, 51). In E. coli., targeting the activation of the stress response pathway in response to ciprofloxacin reduces the evolution of resistant mutant (52). Our findings imply that therapeutic targeting of mutational processes should be beneficial when cancers are most responsive to cytotoxic and targeted therapies while becoming counterproductive once resistance has evolved (28).

Like any modeling study, our simulations represent a simplification of reality. Our analyses of the adaptive state of the populations are limited by the lack of precise knowledge about the underlying Distribution of Fitness Effects (DEFs). Different types of mutational processes can have vastly different DEFs, which was not considered in our simulations. For example, compared to nucleotide level substitutions and small insert/deletions, the effects of large-scale chromosomal aberrations are likely skewed to detrimental effects. Further, the acquisition of therapy resistance to targeted therapies might be dominated by plasticity-mediated mechanisms with varying degrees of heritability, as well as the degree of reprogramming/directionality in the expression level changes (35); this non-genetic variability was not considered in our simulations. Importantly, tumors grow in 3D space, while our simulations and experimental validations were done in the 2D space due to computational and experimental feasibility. Moreover, our choice of on-lattice rather than off-lattice modeling is sub-optimal for accounting for complex interactions and cell motility. Our simulations were performed on a relatively small grid, with a relatively high mutational frequency. Whereas studies with larger population sizes (Supp. Fig. 10) indicated scalability, we cannot exclude the possibility of significant non-linearity at extended grid sizes. Our study did not consider the epistatic interactions between individual mutations, spatial habitats with different phenotypic optima, evolving mutation rates (53), time-dependent selection pressures such as pulsed therapies (54), or interactions between distinct populations. Despite the limitations, our work provides empirical support for an intuitive notion that the adaptive state of a population shapes how mutations affect its evolution. Explicit consideration of the adaptive state of a population can improve the efficiency of therapeutic approaches directed against mutational processes while reducing the potential side effects.

## MATERIALS AND METHODS

### The Agent-Based Model grid definition and agent flowchart

An agent represents a single cell as a pixel in a 2D grid. A grid is an assembly of pixels of a specific size representing a tissue. The model is on-lattice, i.e., cell positions are discretized and correspond to specific pixel coordinates. One agent can occupy at most one pixel and neighborhoods are calculated using Moore approximation. We use a 100 x 100 px domain for all simulations, and each simulation is initialized with 49 cells assembled in a circle at the center of the domain. This ABM is implemented using the HAL library in Java (55) We chose a small grid (100 × 100 px) based on computational efficiency––larger grids and altered mutation rates significantly increase the computational cost (higher mutation rates prolong simulations due to frequent MFA algorithm use, while lower rates require more simulations and longer runtimes).

The flow chart of our ABM, illustrated in Fig. 1a., proceeds as follows: First, a cell dies if a p_1_ < p_die_, where p_1_ ∼U(0,1); dead cells are removed from the domain immediately. If a cell survives, then it can mutate independently of cell division with a probability p_mut_^i^, then divide if p_2_ < p_div_, p_2_ ∼ U(0,1), and if there is available space. If a cell divides, it can mutate again with a probability p_mut_^d^. In this study, we assume p_mut_^i^ = p_mut_^d^ = p_mut_. We restrict p_div_ and p_die_ to p_div_ ∈ [0,1] and p_die_ ∈ [p_die min_,1], where p_die min_ is set to 0.196. This relatively high p_die min_ value ensures a high cell turnover, characteristic of multiple cancer cell lines (29–30, 56). The actions of the ABM are iterated over time and for all alive cells.

To assess the scalability of our results, we compared simulations initialized at 750 and 7500 individuals (7.5% and 75% of carrying capacity) under two DFEs (unbiased, 99%, and 0% neutral mutations). Results (Supp. Fig. 10) indicate that survival near extinction (W_0_ = −0.1, W_0_ = 0) improves with larger populations due to reduced demographic stochasticity (57), but average fitness dynamics remain unchanged.

### Mutation Fitness Algorithm (MFA) and the Distribution of Fitness Effects (DFE)

The MFA algorithm models the mutation-caused quantifiable fitness change in an individual cell: every time a cell acquires a mutation, it impacts cell fitness by changing p_div_ or p_die_. How much fitness changes depends on the underlying DFE, i.e., the frequency of the positive (beneficial), negative (deleterious), or neutral fitness effects. Our DFE is a β(a,b) distribution for all simulations. For results associated with Fig. 2, we set a = b = 3 and define the fitness effects of a mutation x_t_ = X_t_ ∈ [0,1] as negative if x_t_ ∈ [0,x_l_], neutral if x_t_ ∈ ]x_l_, x_u_[, and positive if x_t_ ∈ [x_u_,1]; here, x_l_ = 0.082829 and x_u_ = 0.917171 following (15). We vary a, b, x_l_, and x_u_ when investigating the interplay between the mutation rates, DFEs, and the position on the adaptive landscape (Fig. 3). For all unbiased DFEs (equal frequency of negative and positive mutations), a = b = 3. For the “Major bias” (prevalence of highly impactful negative mutations), a = 1, b = 3, and for the “Minor bias” (prevalence of negative mutations with a variety of impact): a = 1, b = 1.3. We match the specific positive/negative/neutral effect frequencies by solving the cumulative probability distributions for x_l_, and x_u_ values (Supplementary Material 1).

After sampling x_t_, how much p_div_ or p_die_ will change is determined by a linear transformation of x_t_ to y_t_. For negative fitness effects, L^−^≔ [0, x_l_] → [0, 1], L^−^(x_t_) ≔ (x_l_ - x_t_) / x_l_, and for the positive effects L^+^: [x_u_, 1] → [0, 1], L^+^(x_t_) = (x_t_ - x_u_) / (1 - x_u_); for neutral fitness effects y_t_ = 0. Because L^+^ and L^−^ are linear, they are frequency preserving, implying that mutations with smaller impacts happen more frequently than those with large ones (if the DFE is symmetric). Finally, y_t_ can affect p_div_ or p_die_ with a 50% frequency; this corresponds to the p_div_ = p_div_ - y_t_ (50%) or p_die_ = p_die_ + y_t_ (50%) with y_t_ = L^−^(x_t_) (negative effects), and p_die_ = p_die_ - y_t_ (50%) or p_div_ = p_div_ + y_t_ (50%) with y_t_ = L^+^(x_t_) (positive effects). If p_div_ +/- y_t_ exceeds 0 or 1, and p_die_ +/- y_t_ exceeds p_die_min_ (0.196) or 1, the values are set to 0, 1 or p_die_min,_ respectively.

While the p_div_ and p_die_ changes happen immediately, they might be effective first during the next iteration; see supplementary material 1 for the relevant mathematical details. In addition to fitness changes, cells accumulate all mutations, allowing us to track the number of positive, negative, and neutral mutations over time.

### Data collection

Each run of the ABM allows the collection of the average cell data (see supplementary figure 1a). The average cell data (e.g., mean p_div_) allows us to infer the general population behavior over long time scales quickly and is collected every 100^th^ time step until time step 40000, where 1 time step = ½ of a day = 1/730 of a year. Data from each replicate is saved in csv format. Simulation replicates inferred with this approach are denoted by the hat symbol 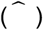. Some replicates can go extinct because of stochasticity, space limitations, and the W_0_. Thus, the total number of analyzed replicates 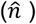 often differs from the total simulation replicates executed (*n*).

### Determining the no. simulations (statistical significance)

We estimated the number of required simulations by capturing the variance in the final number of cells (i.e., at time 54 years = 40000-time steps) as a function of the number of simulation replicates (Supp. Fig. 4) and observed for p_mut_ > 0, the variance converges; however, the asymptotic value of variance and the speed of convergence depend on the p_mut_: higher p_mut_ leads to a higher variance value yet faster variance stabilization. In addition to the p_mut_, the value of stabilized variance depends on the DFE (Supp. Fig. 4). Hence, the apriori determination of the sufficient number of simulations for in-depth statistical analysis is limited. Because populations are more prone to extinction with decreasing W_0_, determining a sufficient number of simulations with low W_0_ can induce a bias; thus, we use W_0_ = 0.75 (initial p_div_ = 1, p_die_ = 0.2) to determine sufficient simulations as n = 250 for p_mut_ = 0, 1e-4 and 1e-3, n = 150 for p_mut_ = 1e-2, n = 75 for p_mut_ = 1e-1, and n = 35 for p_mut_ = 1.

### Max 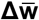

We compute the Max 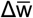 as follows: 1) we first calculate the average fitness over time 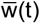 of a population with equal p_mut_ and W_0_, then 2) we compute the forward differences, i.e., 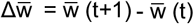, as well as their absolute values, 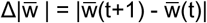; while 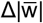 is strictly positive 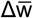 is not, and can thus indicate population fitness increase or decline. In 3), we select the maximum of the 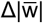, (Max 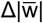) and report the equivalent 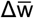 value; this value corresponds to Max 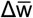. Fitness averages of populations that faced extinctions for all simulations were excluded from this analysis. Because 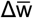 is a numerical approximation calculated from the simulation data, Max 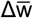 depends on the number of data points and sampling size used for this calculation. To ensure that Max 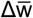 is a representative measure of adaptiveness (i.e., that a significantly larger than zero value of Max 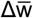 exists), we analyzed the individual 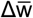 curves; examples of these curves (Supp. Fig. 3a) show that our data collection time step is sufficient to use Max 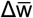 for further analysis.

### Experimental validation studies

ALK+ non-small cell lung cancer cell line H3122 was used for the *in vitro* monolayer assays. The cell line was maintained as a 2D cell model in RPMI media (Fisher), supplemented with 10% fetal bovine serum (FBS) (Fisher) and insulin (ThermoScientific). Mutations were triggered using Ethyl methanesulfonate (EMS) (Fisher). The sensitivity of H3122 cells with the addition of EMS (1µM) in long-term colony formation assay was assessed with DMSO (baseline) and ALKi, Crizotinib (0.5µM), Alectinib (0.5µM) and Lorlatinib (2µM). Briefly, cells were seeded at 0.6 million cells per 6 cm plate and treated with ALKi or ALKi + EMS and, at 2,000 cells per 6 cm plate, treated with DMSO and EMS. For the EMS groups, cells were pretreated with EMS for 48 hours to facilitate mutagenesis before seeding for the assay. Treatment was repeated with fresh media twice a week. Crystal violet staining was performed after 17 days for baseline groups, 36 days for Alectinib groups, 26 days for Lorlatinib groups and 19 days for Crizotinib groups. Images were taken and analyzed using ImageJ (Fiji). Colony numbers, total area covered by the cells and areas of each colony were quantified. The clonogenic fraction was calculated as the fraction of the number of colonies by cells plated. 3 independent experiments were performed. The short-term sensitivity of cells was assessed via cell titer glo assay. Briefly, 4,000 H3122 cells were seeded per well in a 96 well plate and treated with DMSO or ALKi, alone or in combination with EMS at the same concentrations as above. Cell titer glo was carried out after 48 hours of treatment, and % of viability was calculated by normalizing the luminescence of each condition to the DMSO control. All graphs were prepared in GraphPad Prism. Since we have not observed a lot of movement in the cell cultures, we have not conducted a cell migration assay.

## Notes

https://github.com/gortzaah/AdaptationMutation

### Competing Interest Statement

The authors have declared no competing interest.

### Summary of Updates

The revised manuscript contains more detailed description of the experimental work and some figures were redesigned for clarity

