## Supplementary material for "The adaptive state determines the impact of mutations on evolving populations"

PNAS

##### This PDF file includes:

Supporting text
Figs. S1 to S13
Legends for Movies S1 to S18

##### Other supporting materials for this manuscript include the following:

Movies S1 to S18

### Supporting Information Text

#### 1. Supplementary figures

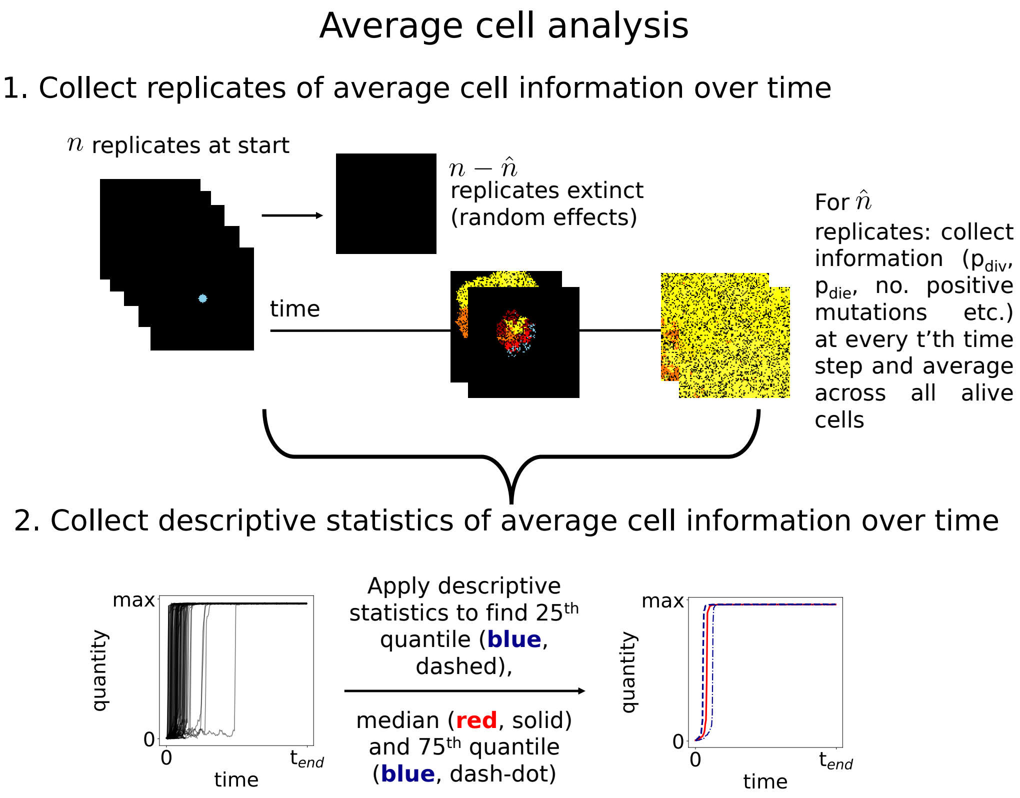


**Fig. S1** – Average cell data sampling and processing pipeline. 1) Data collection for statistical analysis. Typical cell data values collected are time, "no_cells" (no. cells), "pdiv" (probability of division), "pdie" (probability of death), "pos_muts" (no. positive mutations), "neutral_muts" (no. neutral mutations), "neg_muts" (no. negative mutations), "pos_effect" (positive effect of mutation), "neg_effect" (negative effect of mutations), "no_pro" (no. of dividing cells), and "pspace" (the probability of having free space for cell division). The average positive and negative effects of mutations on the cells were discarded from any analysis performed in this paper due to bad algorithm implementation. 2) The average cell data $\hat{n}$ replicates are statistically analyzed over time by descriptive statistics methods. The default sampling time is the 100th time step.

**Supplementary Figure 1: ABM data collection and processing.**


Malgorzata Tyczynska Weh, Pragya Kumar, Viktoriya Marusyk, Andriy Marusyk, David Basanta

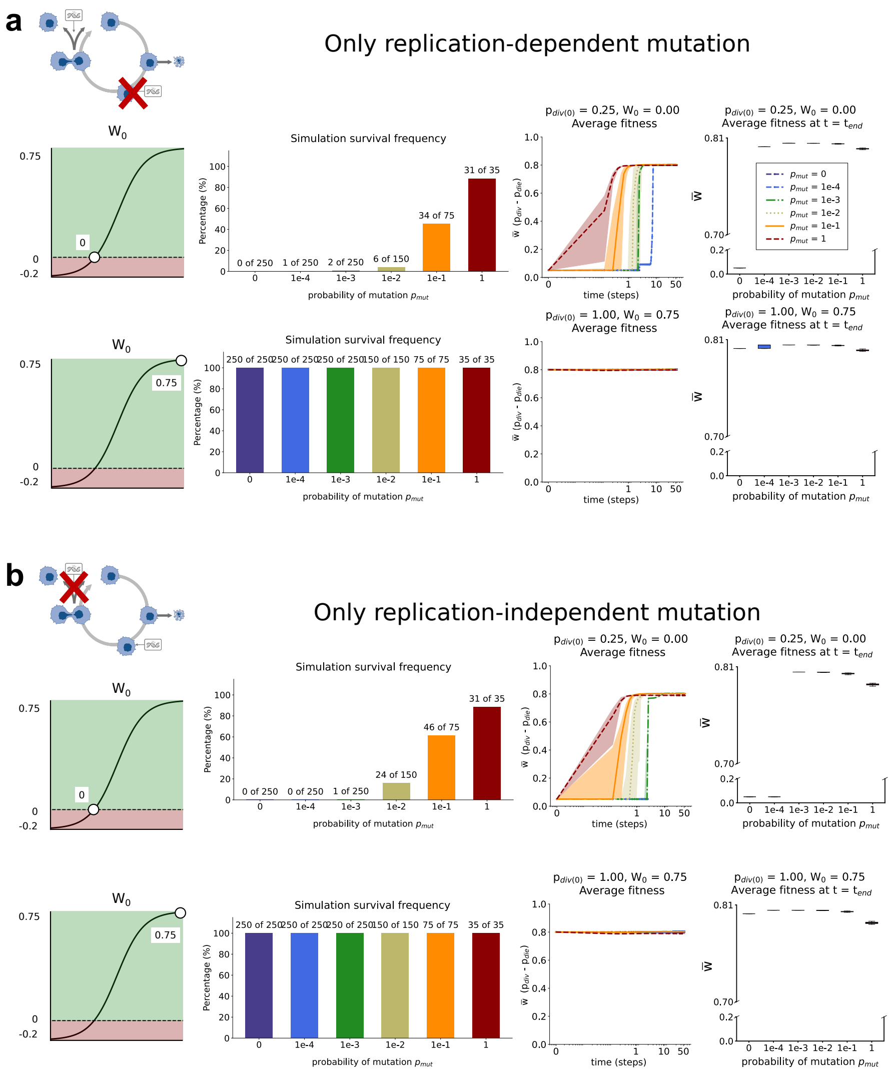


**Fig. S2** – **a:** $p^i_{mut} > 0, p^d_{mut} = 0$, **b**: $p^d_{mut} > 0, p^i_{mut} = 0$. For each panel, we capture the average number of cells (left), average fitness (middle), and the chances of survival (right), with n = 250 ($p_{mut}$ = 0, 1e-4, 1e-3), n = 150 ($p_{mut}$ = 1e-2), n = 75 ($p_{mut}$ = 1e-1), and n = 35 ($p_{mut}$ = 1) simulations. All simulations are initialized with $W_0$ = 0 (initial $p_{div}$ = 0.25, $p_{die}$ = 0.2) and with 49 cells.

**Supplementary Figure 2: The impact of the absence of division-independent (a) or division-dependent mutation (b) at the bottom ($W_0$ = 0) or peak ($W_0$ = 0.75) of the adaptive landscape..**

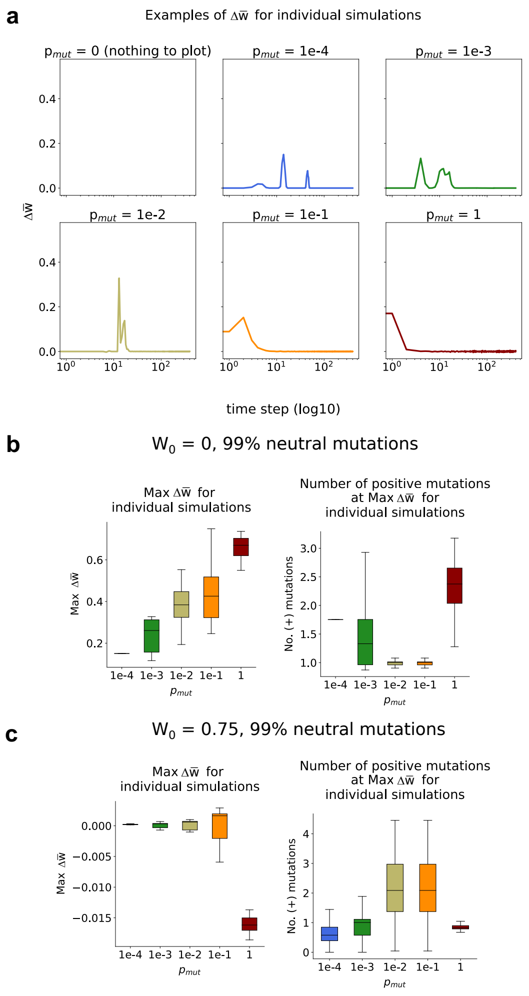


**Fig. S3** – Max $\Delta$w. **a**: examples of individual Max $\Delta$w trajectories for conditions as in main figure 2. **b,c**: Max $\Delta$w values of individual trajectories for conditions as in the main figure 2 (**b**) and at the top of adaptive landscape (**c**), indicated by $W_0$ = 0.75.

**Supplementary Figure 3: Max $\Delta w$ for individual simulations and the corresponding no. of positive mutations.**

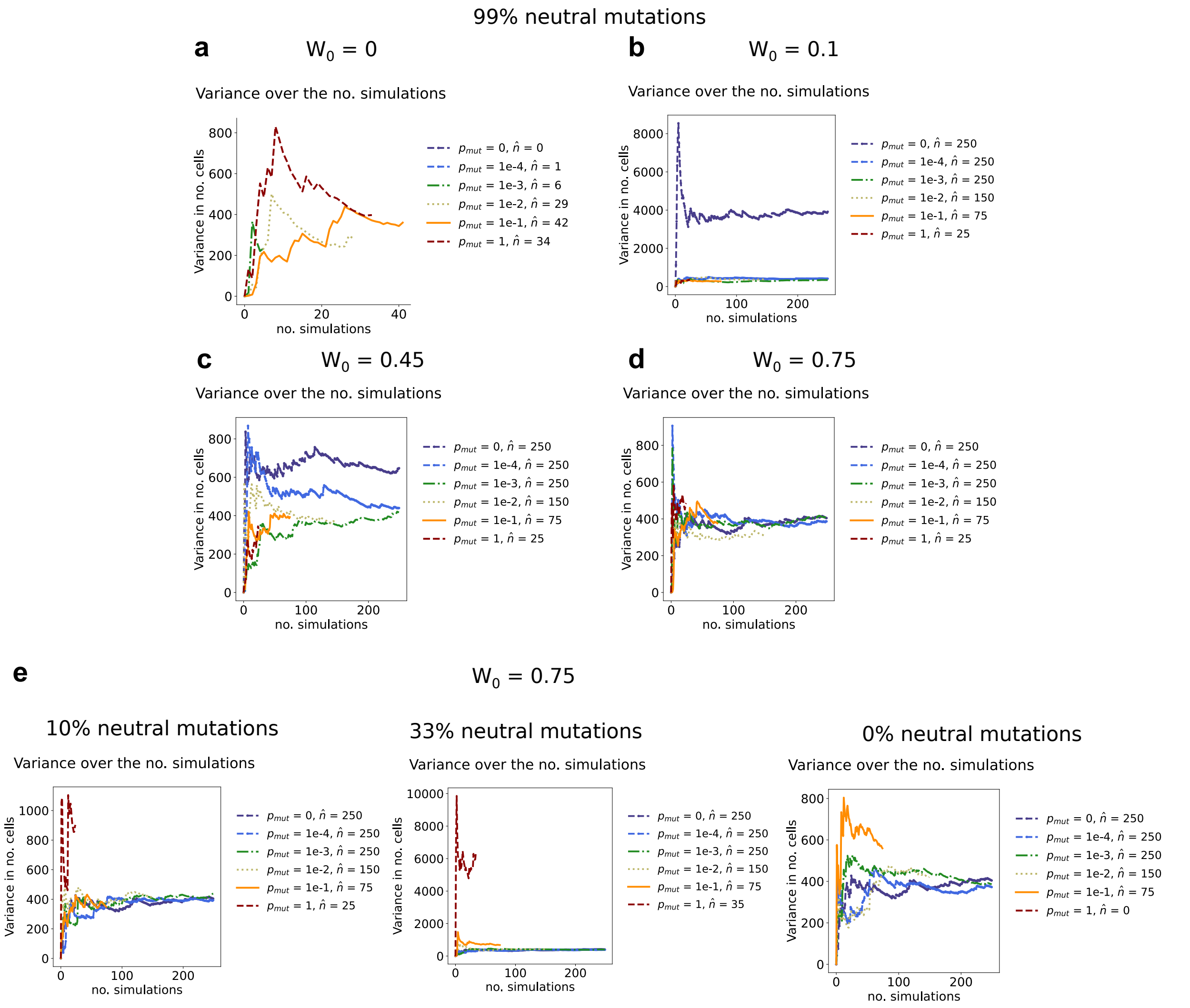


**Fig. S4** – The sample variance in the no. cells at the last time point as a function of the no. simulation replicates. The last time point corresponds to 54 years = 40000 time steps For a-**d**, the DFE is 99% neutral mutations, 0.5% positive and negative mutations (as in Main Figure 2), and $W_0$ = 0 (**a**), $W_0$ = 0.1 (**b**), $W_0$ = 0.45 (**c**), and $W_0$ = 0.75 (**d**). The DFE changes in **e** to have 10% neutral mutations (bottom left), 33% neutral mutations (bottom middle), and 0% neutral mutations (bottom right), but $W_0$ = 0.75. For all figures, $\hat{n}$ describes the number of simulations that survived out of n = 250 ($p_m ut$ = 0, 1e-4, 1e-3), n = 150 ($p_m ut$ = 1e-2), n = 75 ($p_m ut$ = 1e-1), and n = 35 ($p_m ut$ = 1).

**Supplementary Figure 4: Example of variance in the number of cells as a function of the number of simulations .**

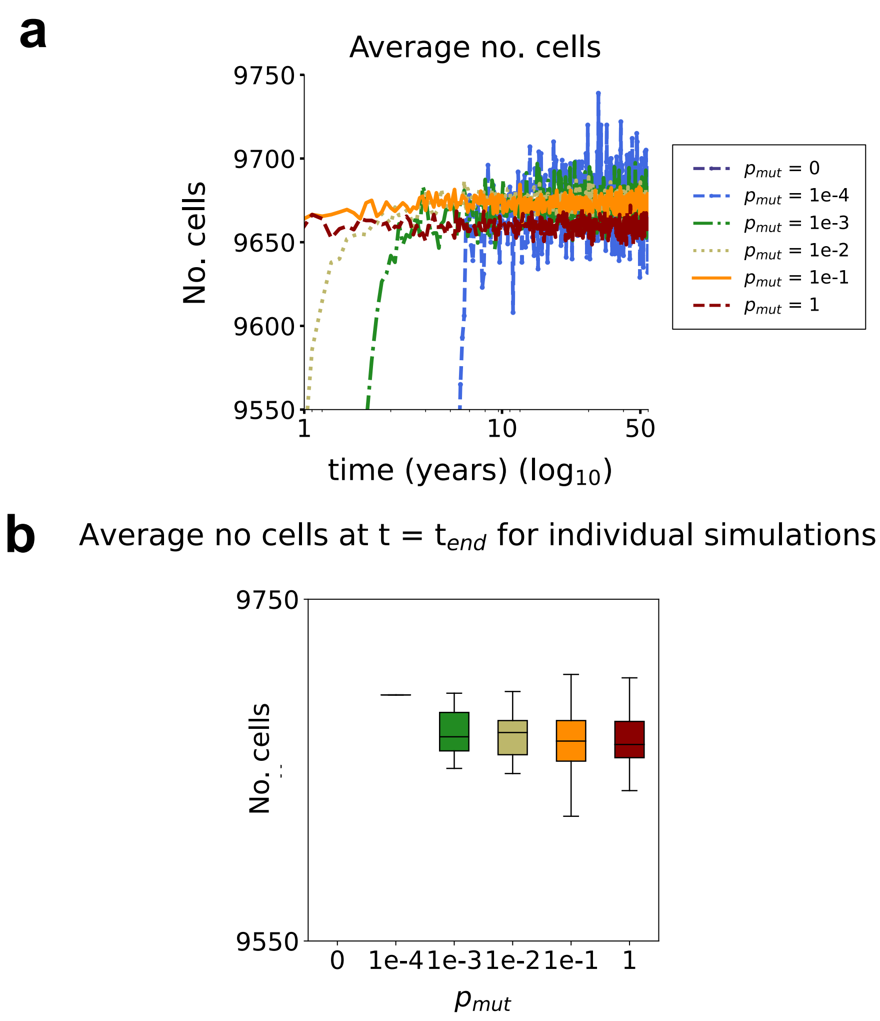


**Fig. S5** – Zoom-in on the average number of cells (**a**) and the boxplots of the average number of cells at the last time point of from the individual simulations (**b**). Simulation parameters follow the description of Fig. 2.

**Supplementary Figure 5: No. cells at t = t$_{e}nd$ (Support to Main Fig. 2) .**

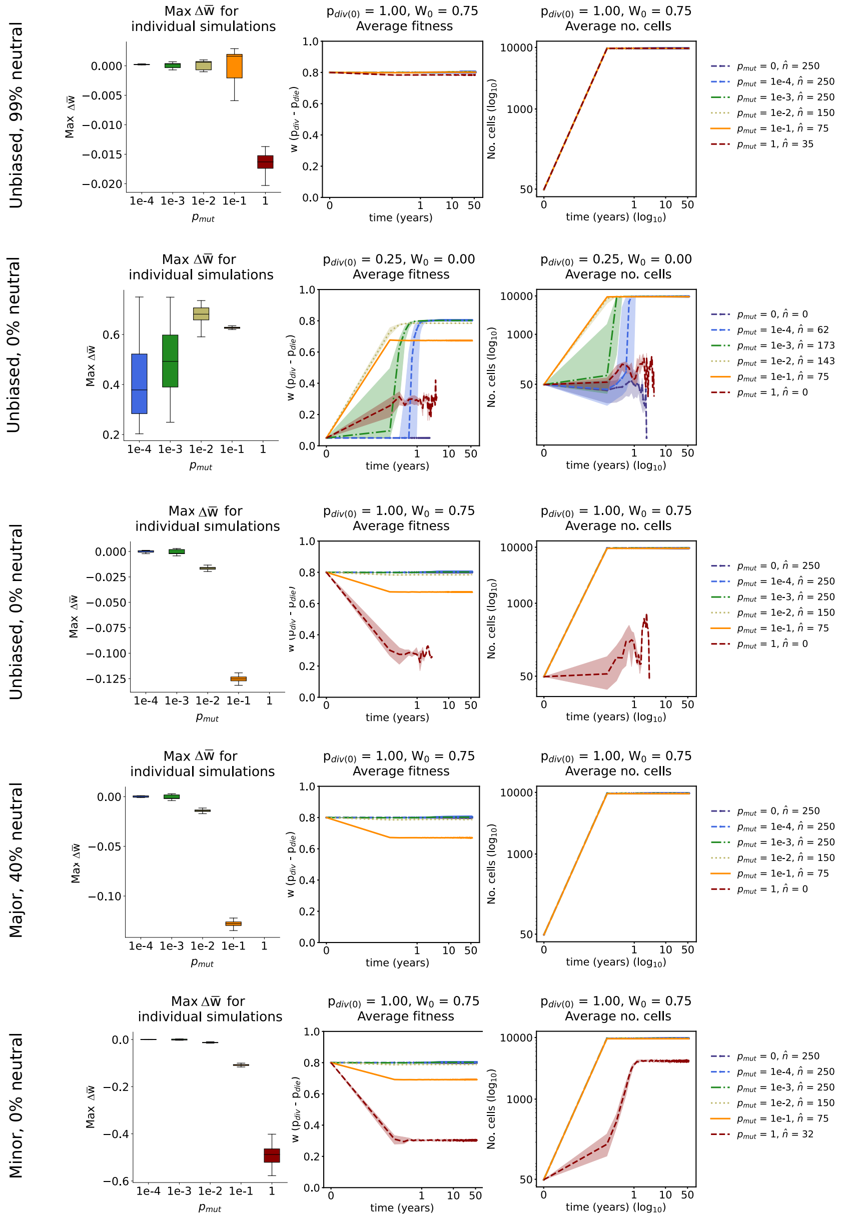


**Fig. S6** – Example of Max $\triangle$w, average fitness trajectories, and the population size. The description of colors and variables follows from Main Fig. 2.

**Supplementary Figure 6:Illustrative examples of Max $\triangle$w, average fitness trajectories, and the population size (Support to Main Fig. 3) .**

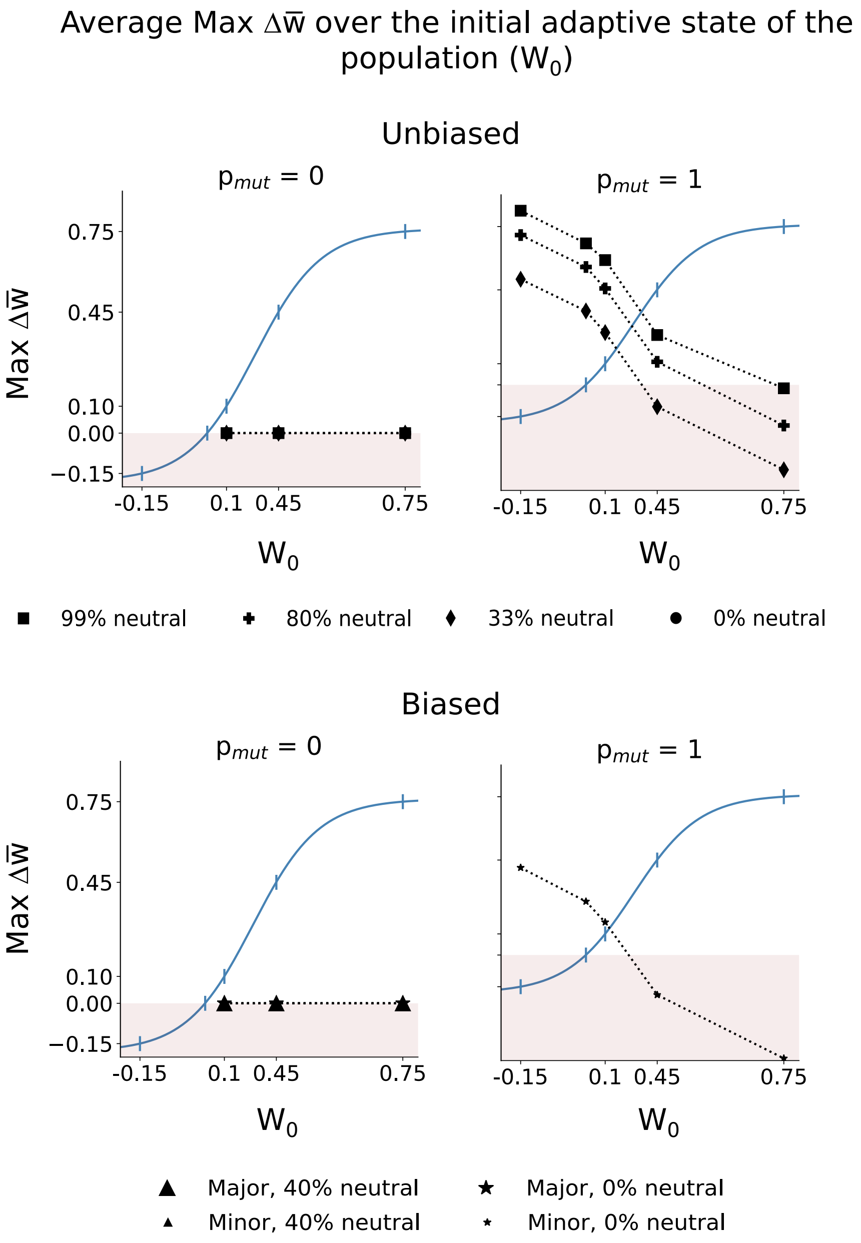


**Fig. S7** – Max $\Delta\bar{w}$ over the adaptive landscape for $p_{mut}$ = 0 (left column) and $p_{mut}$ = 1 (right column); conditions and descriptions follow Main Fig. 3b.

23 **Supplementary Figure 7: Max $\Delta\bar{w}$ over the adaptive landscape for $p_{mut}$ = 0 and $p_{mut}$ = 1 (Support to Main Fig. 3).**

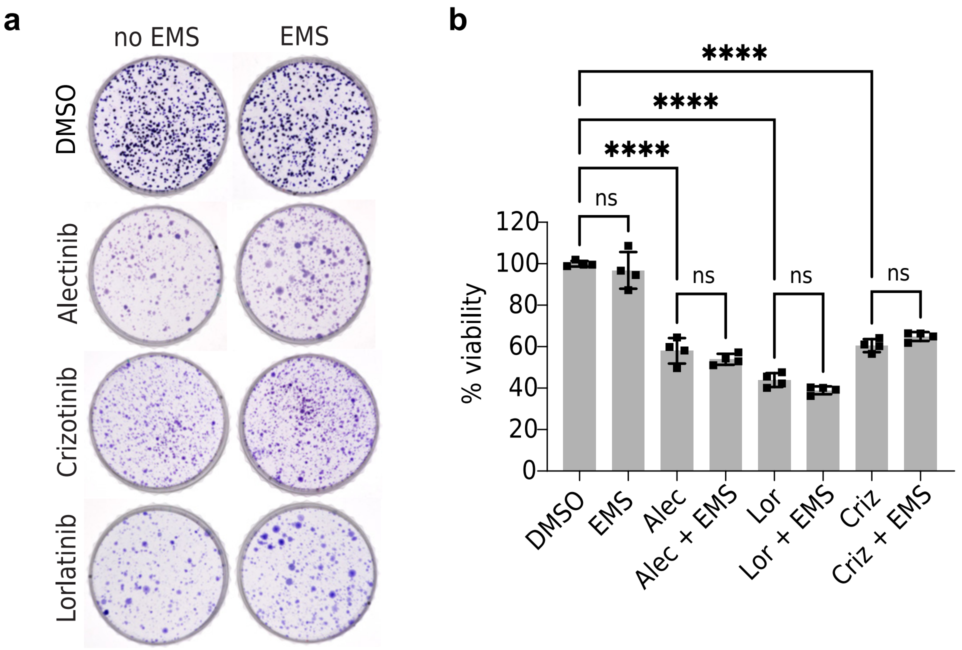


**Fig. S8** – **a**: Representative images of colonies of H3122 cells treated with DMSO or ALKi, Alectinib (0.5$\mu$M), Crizotinib (0.5$\mu$M) or Lorlatinib (2$\mu$M), alone or in combination with EMS (1$\mu$M). **b**:Cell Titer Glo viability assay of H3122 cells treated with DMSO, Alectinib (0.5$\mu M$), Crizotinib (0.5$\mu M$) or Lorlatinib (2$\mu M$), alone or in combination with EMS (1$\mu M$) for 48 hours. The significance testing was done with the Mann-Whitey test; p-values – <0.05 (*), <0.01 (**), <0.001 (***), <0.0001 (****).

**Supplementary Figure 8: Cell cultures.**

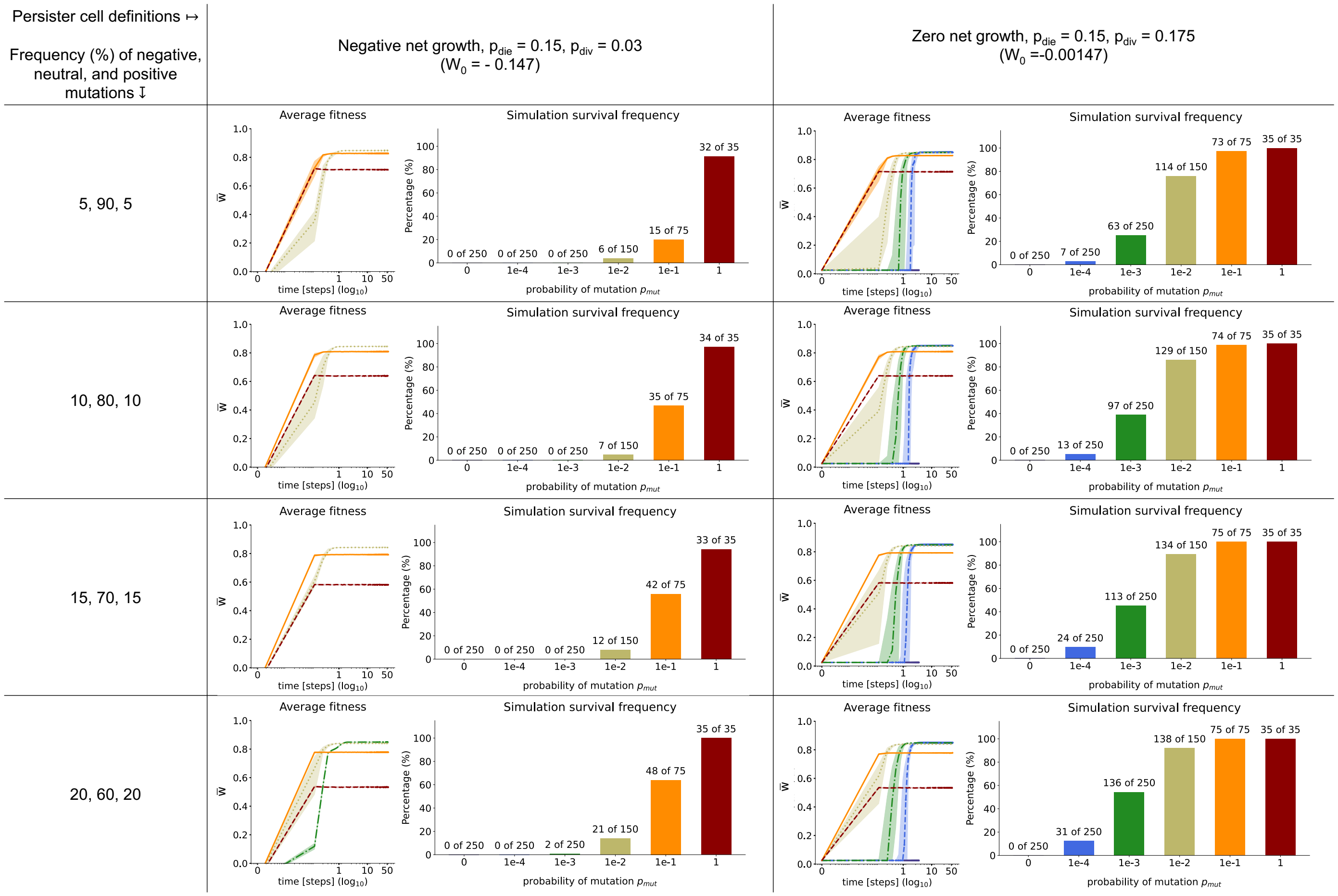


**Fig. S9** – Average population fitness and the simulation survival frequency conducted for two alternative definitions of the persister cell state and different frequencies of positive, negative and neutral mutations. The second column represents the definition of persister cell states where population net growth rate is negative ($p_{div} = 0.03$), and the third: populations with zero net growth rate but lower proliferation probability ($p_{div} = 0.175$; for both cases, $p_{die} = 0.15$. For all cases, the minimum cell death probability is $p_{die,min}$ is $p_{die} = 0.15$, and the mutation distribution is $\beta(3, 3)$.

#### Supplementary Figure 9: Changing the persister cell state definition.

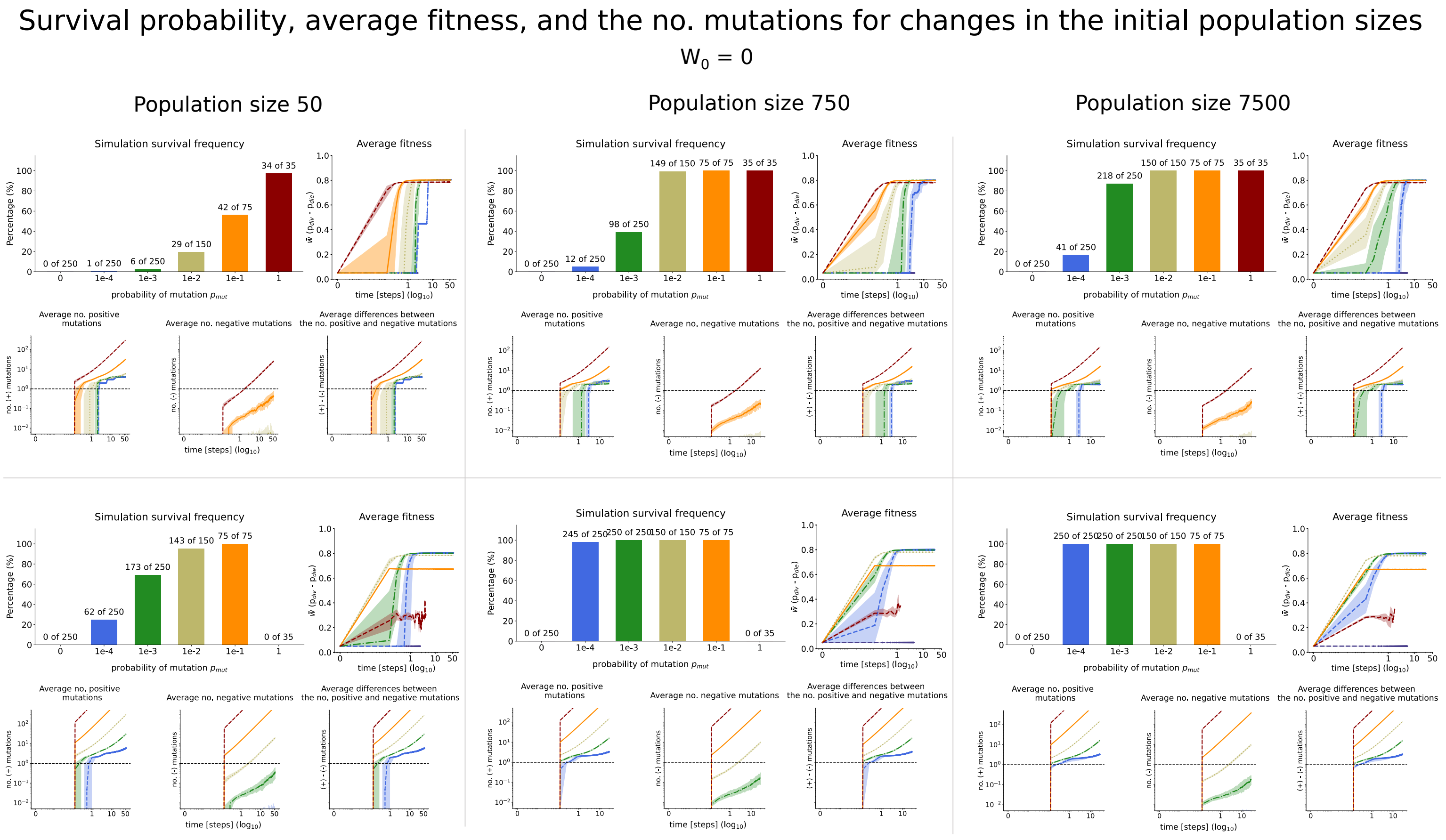


**Fig. S10** – Survival simulation frequency, average population fitness and the average number of mutations per cell for altered initial population sizes, and the unbiased DFE ($\beta(3,3)$) with either 99% or 0% of neutral mutations at $W_0 = 0$ with $p_{div} = 0.25$, $p_{die} = 0.2$, $p_{die,min} = 0.196$. The carrying capacity is kept constant at 10000 cells.

#### Supplementary Figure 10: Changing the initial number of cells.

#### 2. Supplementary material 1: Mathematical modeling of the mutation impacts on cell fitness

**A. Mutation-Fitness (MF) algorithm and bounds.** Every Distribution of Fitness Effects (DFE) is assumed to be a $\beta(a,b)$ distribution; $a = b$ results in symmetric distributions and $a \neq b$ - asymmetric, see fig S11. $\beta(a,b)$ are compact on the $[0,1]$ interval, thus suitable for modeling probability distributions of fitness effects. When investigating the interplay between DFEs, position on the adaptive landscape,

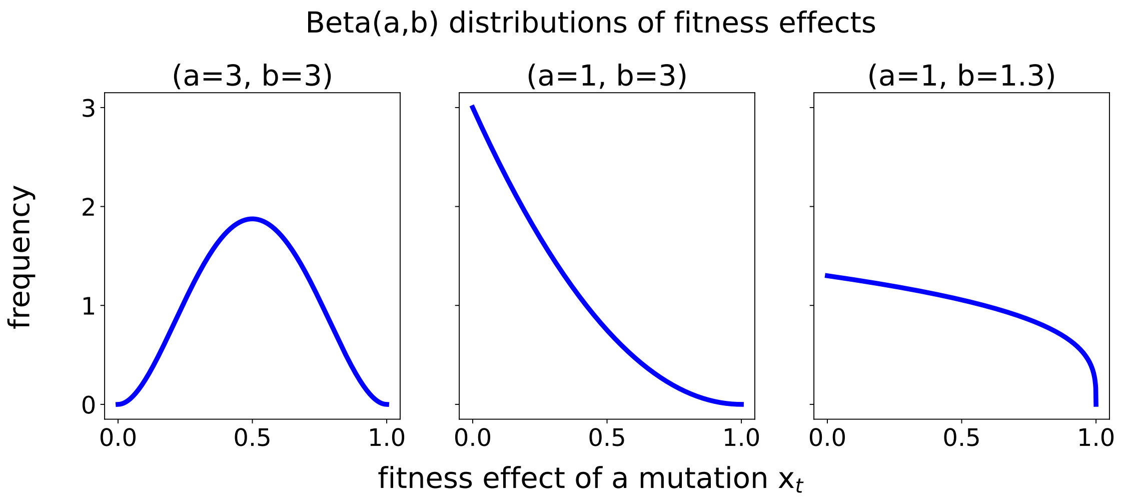


**Fig. S11** – The $\beta(a,b)$ distribution used to model different DFEs.

and the mutation rates, we use $\beta(3,3)$ (fig S11 left) to model all symmetric the DFEs, the $\beta(1,3)$ (fig S11 middle) to model the "major" bias to the negative fitness effects, and $\beta(1,1.3)$ (fig S11 right) to model the minor bias of DFEs to the negative fitness effects. Regardless of the distribution, the negative fitness effects correspond to $x_t \in [0, x_l]$, neutral to $x_t \in ]0, x_u[$ and $x_t \in [x_u, 1]$. The values $x_l$, $x_u$ are solutions to the equations of the cumulative distribution function corresponding to the specific frequencies of mutations. For example, with $\beta(3,3)$ and 99% of neutral mutations, we have 0.5% of negative and positive mutations each, and thus must solve the following two equations for $x_l$ and $x_u$:

$$\int_0^{x_l} \beta(3,3)(x_t)\mathrm{d}x_t = 0.005 \Rightarrow x_l = 0.082829$$

$$\int_{x_u}^{1} \beta(3,3)(x_t)\mathrm{d}x_t = 0.005 \Rightarrow x_u = 0.917171$$

The following table shows all the values for $x_l$, $x_u$ and the corresponding frequency of fitness effects for the symmetric distributions ($\beta(3,3)$):

| % neutral | $x_l$ | $x_u$ |
|---|---|---|
| 99 | 0.082829 | 0.917171 |
| 80 | 0.246636 | 0.753364 |
| 33 | 0.409128 | 0.590872 |
| 0 | 0.5 | 0.5 |

The following table shows all the values for $x_l$, $x_u$ and the corresponding frequency of fitness effects for the distribution ($\beta(1,3)$) with major skew towards negative fitness effects:

| % negative; % neutral; % positive | $x_l$ | $x_u$ |
|---|---|---|
| 40;0;60 | 0.1565673 | 0.1565673 |
| 40,40,20 | 0.1565673 | 0.4151964 |

Finally, this table shows all the values for $x_l$, $x_u$ and the corresponding frequency of fitness effects for the distribution ($\beta(1,1.3)$) with minor skew towards negative fitness effects:

| % negative; % neutral; % positive | $x_l$ | $x_u$ |
|---|---|---|
| 40;0;60 | 0.32493263 | 0.32493263 |
| 40,40,20 | 0.32493263 | 0.7100449 |

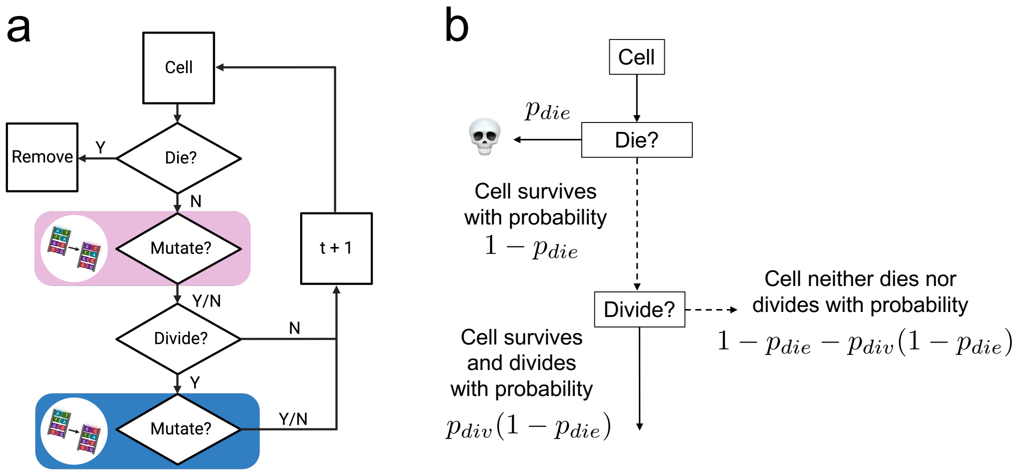


**Fig. S12** – a: ABM schemata as in the main text. b: Simplified ABM schemata illustrating conditional probabilities used in example 2.1. The solid arrows symbolize "yes" answers, the dashed ones: "no".

**B. Introduction to the ABM and conditional probabilities.** Given the ABM with structure as in fig. S12:
The cell-fate decisions at time $t$ are executed based on the flowchart with boxes arranged in a specific order. Each box stands for a specific decision, e.g., the first box is about death: when the cell is at the box "die?", it can die with probability $p_{die}$ and survive with the probability $(1 - p_{die})$; formally, with a $p \sim U(0, 1)$, a cell dies if $p \leq p_{die}$. When the cell is at the box "Mutate?", it mutates independently of cell division with a probability $p^i_{mut}$; when it is at "Division?", the division is executed with probability $p_{div}$; then if a cell is at second mutation box, it divides with division-dependent probability $p^d_{mut}$. Because all $p_{die}$, $p_{div}$, $p^i_{mut}$ and $p^d_{mut}$ are determining the outcome at each box, we call them the *effective* probabilities. Note that each of these probabilities is time-dependent, i.e., $p_{die} := p_{die}(t)$, but the dependency on $t$ is here omitted for the sake of readability.
Nevertheless, the paths to reach each decision box (e.g. division) are not independent of other actions: for example, a cell must be alive to divide. To fully understand the mathematical consequences of this statement, consider the following example (note, that we omit the space limitations for the sake of clarity):

**Example 2.1** *Assume $p^i_{mut} = p^d_{mut} = 0$, implying that $p_{div}$ nor $p_{die}$ are not altered by mutations. From fig S12b, we observe that the reproductive success of a cell depends on its ability to survive – (i.e., only alive cells can divide). Mathematically, this implies that the true division probability of a cell ($\hat{p}_{div}$) is conditional on its survival, i.e., $\hat{p}_{div} = p(p_{div} \| (1 - p_{die}))$. Because $p_{div}$ and $p_{die}$ are independently assigned, we have $\hat{p}_{div} = p_{div} \times (1 - p_{die})$. Consequently, at time $t$, a cell has a probability of not dying and not dividing equal to $1 - p_{die} - \hat{p_{div}} = 1 - p_{die} - p_{div}(1 - p_{div})$. These probabilities sum to one, i.e. $p_{die} + p_{div}(1 - p_{die} + (1 - p_{die} - p_{div}(1 - p_{div})) = 1$.*

**C. The effects of mutations on the consistency conditional probabilities.** The point of the previous section is that conditional probability theory is essential to interpret the ABM's logic - how individual cell dynamics lead to the emergent population patterns - correctly. Nevertheless, in the example 2.1, we assumed $p^i_{mut} = p^d_{mut} = 0$. How do the non-zero values of $p^i_{mut}$ $p^d_{mut}$, i.e. the potential change of the values of $p_{div}$ and $p_{die}$, alter this analysis?
Let $\Lambda$ be the permutation of the probability $p_{div}$ or $p_{die}$ describing the effect of the mutation on one of these parameters at time $t$ (see Fig. 1b in the main text), i.e. when a mutation happens, $p_{(.)}$ takes value $\Lambda(p_{(.)})$.
Let $p^i_{mut} > 0$ and $p^d_{mut} = 0$. The cell can potentially die with probability $p_{die}$ or survive with probability $(1 - p_{die})$. Then, assume that a division-independent mutation happens with a non-zero effect on $p_{die}$. Because the decision about death or survival is made upon "Die?" box, not upon "Mutate?" box, a cell still survives with probability $(1 - p_{die})$, and not with $\Lambda(p_{die})$. Consequently, the cell divides with the probability $p_{div}(1 - p_{die})$ and neither dies nor divides with probability $1 - p_{die} - p_{div}(1 - pdie)$. See fig. S13a for the illustration.
A division-independent mutation affecting $p_{div}$ changes the $p_{div}$ before a cell decides to divide. Hence, the mutation causes division conditioned upon survival with probability $\Lambda(p_{div})(1 - p_{die})$. The cell hence neither dies nor divides with the probability $1 - p_{die} - \Lambda(p_{div})(1 - p_{die})$. This is illustrated on fig. S13b.
Consider now $p^i_{mut} > 0$ and $p^d_{mut} > 0$ and figure S12a. Even if $\Lambda$ affects either $p_{die}$ or $p_{div}$, the algorithm proceeds directly to the next time iteration. Hence, regardless of whatever changes are made for the values $p_{div}$ or $p_{die}$, they are first used in the next time iteration.

**D. Derivation of zero net growth conditions.** Initially, all cells in the ABM are initialized with no mutations and with equal $p_{die}$ and $p_{div}$ for all cells. From example 2.1, the cells have equal conditional division and death rates if:

$$p_{div}(1 - p_{die}) = p_{die},$$

implying that if all cells are having identical $p_{die}$ and $p_{div}$, then the populations grow at a rate:

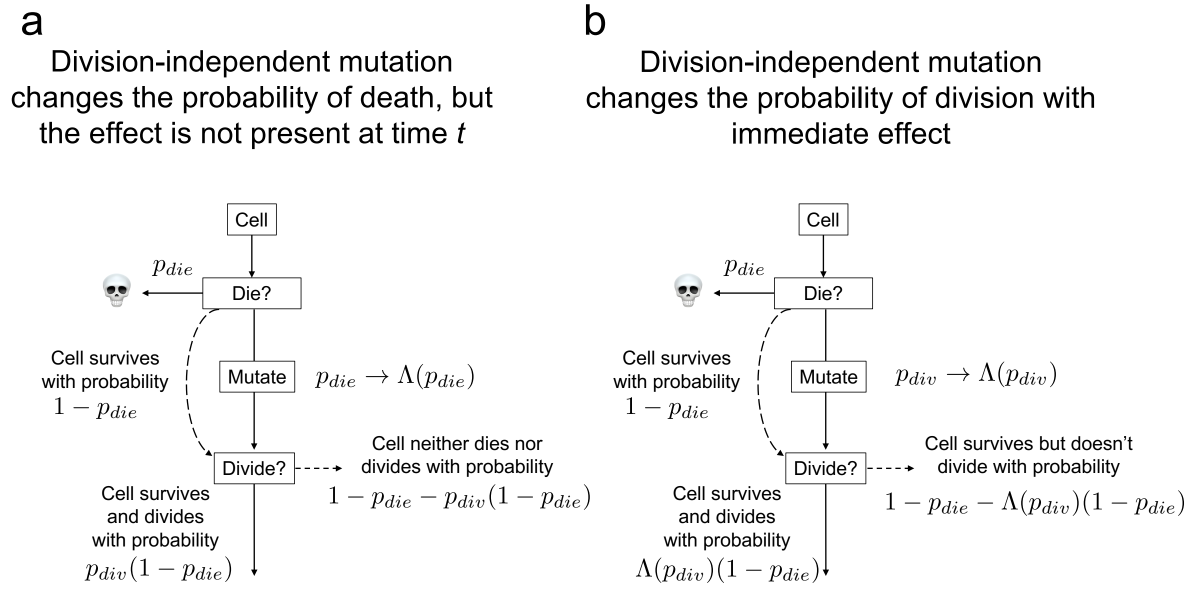


**Fig. S13** – The effect of the division-independent mutation affecting either $p_{die}$(left). The solid arrows symbolize "yes" answers, the dashed ones: "no".

$$r = p_{div}(1 - p_{die}) - p_{die} \Leftrightarrow \frac{1}{(1 - p_{die})} r = p_{div} - \frac{p_{die}}{(1 - p_{die})},$$

implying that populations have zero net growth ($r = 0$) if:

$$p_{div} - \frac{p_{die}}{(1 - p_{die})} = 0.$$

Consequently,

$$W_0 = p_{div} - \frac{p_{die}}{1 - p_{die}}, \qquad [1]$$

corresponds to the population net growth rate **if all individuals are identical**: if $W_0 < 0$, then $p_{div} < \dfrac{p_{die}}{1 - p_{die}}$ and populations are expected to extinct. Conversely, if $W_0 > 0$, then $p_{div} > \dfrac{p_{die}}{1 - p_{die}}$ resulting in population growth. Because during the simulation, cells acquire mutations that cause changes in $p_{div}$ and $p_{die}$, the $W_0$ is a valid representation of the population dynamics only at the beginning, when all cells are identical. While the assumption about identical individuals is essential for $W_0$'s validity, this measure is imperfect: stochasticity and spatial effects can cause fluctuations in cell population numbers.

**E. Implications for division-dependent and division-independent mutations.** While the previous sections focused on the $p_{div}$ and $p_{die}$, we still do not understand how either division-independent or division-dependent mutations can impact population dynamics, causing adaptation. With $p^i_{mut}$ and $p^d_{mut}$ describing the division-independent and division-dependent mutation rates, respectively, we can derive the conditional probabilities driving their impact on cell dynamics leading to emergent patterns on the population level using Fig S12a and conditional probabilities, as follows:

$$\hat{p}^i_{mut} = p^i_{mut}(1 - p_{die}),$$

thus, a division-independent mutation depends on the cell's survival. Now, recall that the mean initial value of $p_{die}(0)$ is $0.2$ for most of the analysis and that the minimal allowed value of the probability of death, $p_{die,min}$, is $0.196$. Because we observe $p_{die} \rightarrow p_{die,min}$ during adaptation (see the supplementary movie set 3), we must have:

$$p^i_{mut}(1 - p_{die}) \rightarrow p^i_{mut}(1 - p_{die,min}) \Leftrightarrow 0.8 p^i_{mut} \rightarrow 0.804 p^i_{mut} \text{ as } t \rightarrow \infty, \qquad [2]$$

thus, the probability of division-independent mutation having an impact during the simulation changes very little.

The division-dependent mutation $p^d_{mut}$ is conditionally dependent on cell survival $(1 - p_{die})$ and cell division $p_{die}$:

$$\hat{p}^d_{mut} = p^d_{mut}(1 - p_{die})p_{div}$$

Given the typical initial condition $p_{div}(0) = 0.25$, we have that the initial division-dependent mutation is scaled with $(1-p_{die}(0))p_{div}(0) = 0.8 \times 0.25 = 0.2$. As cells adapt, they increase their $p_{div}$ towards its maximal value $1$ and decrease $p_{die}$ towards $p_{die,min}$. Consequently:

$$p^d_{mut}(1 - p_{die})p_{div} \rightarrow p^d_{mut}(1 - p_{die,min})p_{div,max} \Leftrightarrow 0.2p^d_{mut} \rightarrow 0.80p^d_{mut} \text{ as } t \rightarrow \infty. \quad [3]$$

Comparing the expressions Eq. (2) with Eq. (3) we observe that the $p^i_{mut}$ has almost the same impact throughout the whole course of adaptation. This is in contrast to Eq. (3), which experiences a large change in factor from $0.204$ to $0.804$. Because of the low initial impact, the division-dependent mutations have a lower chance of occurrence during the early stages of adaptation. This causes more stochastic initial dynamics, as seen in Supplementary Fig 6. On the other hand, this result suggests that division-independent mutations are crucial for adaptation in its earliest phase. Biologically, these mutations can be induced by any form of environmental stress, including the stress from the drug. Finally, note that the impacts of $p^d_{mut}$ and $p^i_{mut}$ on cell dynamics are asymptotically the same, thus over time, thus the difference between the impact of division-dependent and division-independent mutations fades over time.

#### Supplementary Movies

**Movie S1. ABM simulation for the initial probability of division 0.25 and probability of mutation 0, example 1**

Example of the .gif file of the ABM simulations with $p_{mut} = 0$, $p_{div} = 0.25$.

**Movie S2. ABM simulation for the initial probability of division 0.25 and probability of mutation 0, example 2**

example of the .gif file of the ABM simulations with $p_{mut} = 0$, $p_{div} = 0.25$.

**Movie S3. ABM simulation for the initial probability of division 0.25 and probability of mutation 0, example 3**

example of the .gif file of the ABM simulations with $p_{mut} = 0$, $p_{div} = 0.25$.

**Movie S4. ABM simulation for the initial probability of division 0.25 and probability of mutation 1, example 1**

example of the .gif file of the ABM simulations with $p_{mut} = 1$, $p_{div} = 0.25$.

**Movie S5. ABM simulation for the initial probability of division 0.25 and probability of mutation 1, example 2**

example of the .gif file of the ABM simulations with $p_{mut} = 1$, $p_{div} = 0.25$.

**Movie S6. ABM simulation for the initial probability of division 0.25 and probability of mutation 1, example 3**

example of the .gif file of the ABM simulations with $p_{mut} = 1$, $p_{div} = 0.25$.

**Movie S7. Fitness, the number of positive and negative mutations for the probability of mutation 1 the and initial probability of division 0.25 over time**

Each graph is obtained from analyzing single-agent data from 10 simulations collected every 100 time step.

**Movie S8. Fitness, the number of positive and negative mutations for the probability of mutation 1e-1 the and initial probability of division 0.25 over time**

Each graph is obtained from analyzing single-agent data from 15 simulations collected every 100 time step.

**Movie S9. Fitness, the number of positive and negative mutations for the probability of mutation 1e-2 the and initial probability of division 0.25 over time**

Each graph is obtained from analyzing single-agent data from 15 simulations collected every 100 time step.

**Movie S10. Fitness, the number of positive and negative mutations for the probability of mutation 1e-3 the and initial probability of division 0.25 over time**

Each graph is obtained from analyzing single-agent data from 15 simulations collected every 100 time step.

**Movie S11. Fitness, the number of positive and negative mutations for the probability of mutation 1e-4 the and initial probability of division 0.25 over time**

Each graph is obtained from analyzing single-agent data from 15 simulations collected every 100 time step.

**Movie S12. Fitness, the number of positive and negative mutations for the probability of mutation 0 the and initial probability of division 0.25 over time**

Each graph is obtained from analyzing single-agent data from 15 simulations collected every 100 time step.

**Movie S13. Fitness, the number of positive and negative mutations for the probability of mutation 1 the and initial probability of division 1 over time**

Each graph is obtained from analyzing single-agent data from 10 simulations collected every 100 time step.

**Movie S14. Fitness, the number of positive and negative mutations for the probability of mutation 1e-1 the and initial probability of division1 over time**

Each graph is obtained from analyzing single-agent data from 15 simulations collected every 100 time step.

**Movie S15. Fitness, the number of positive and negative mutations for the probability of mutation 1e-2 the and initial probability of division 1 over time**

Each graph is obtained from analyzing single-agent data from 15 simulations collected every 100 time step.

**Movie S16. Fitness, the number of positive and negative mutations for the probability of mutation 1e-3 the and initial probability of division 1 over time**

Each graph is obtained from analyzing single-agent data from 15 simulations collected every 100 time step.

**Movie S17. Fitness, the number of positive and negative mutations for the probability of mutation 1e-4 the and initial probability of division 1 over time**

Each graph is obtained from analyzing single-agent data from 15 simulations collected every 100 time step.

**Movie S18. Fitness, the number of positive and negative mutations for the probability of mutation 0 the and initial probability of division 1 over time**

Each graph is obtained from analyzing single-agent data from 15 simulations collected every 100 time step.
